# FoldMark: Safeguarding Protein Structure Generative Models with Distributional and Evolutionary Watermarking

**DOI:** 10.1101/2024.10.23.619960

**Authors:** Zaixi Zhang, Ruofan Jin, Jiangle Liu, Guangxue Xu, Xiaotong Wang, Marinka Zitnik, Huimin Zhao, Le Cong, Mengdi Wang

## Abstract

Protein generative models are transforming biomolecular design and broad applications in bioengineering, drug discovery, and synthetic biology. However, their rapid progress introduces critical challenges, including dual-use biosecurity risks to create pathogenic proteins or bioweapons, and intellectual-property concerns related to unauthorized model use and redistribution. Here, we introduce *FoldMark*, a first-of-its-kind digital watermarking strategy leveraging *distributional* and *evolutionary* principles tailored for protein generative models. FoldMark pretrains a watermark encoder–decoder pair to subtly modify structures, embedding watermark codes guided by evolutionary signals that reduce noise in conserved regions and use flexible regions to increase capacity, ensuring retrievability without loss of structural quality. FoldMark achieves over 95% watermark bit accuracy at *32 bits* with minimal impact on structural quality (>0.9 scTM scores) for leading models including AlphaFold3, RFDiffusion, and RFDiffusionAA. For user tracing, FoldMark can successfully trace up to *1 million* users. Finally, we applied FoldMark to structure-based design of EGFP, CRISPR-Cas13, DHFR, and TEV enzymes, showing wildtype-level functionality for EGFP and Cas13 (avg. 98% fluorescence and avg. 95% editing efficiency), retained activity in 2 of 3 top-ranked enzyme variants, and >90% watermark detection success rates, demonstrating its practical utility for safeguarding AI-driven protein research.

## Introduction

Proteins are life’s essential building blocks and the basis of all living organisms. Understanding their structure is key to uncovering the mechanisms behind their function. With the advancement of generative AI [*1*], protein structure generative models have revolutionized both protein structure prediction [*2, 3*] and de novo protein design [*4, 5, 6*]. For example, AlphaFold2 [*2*] made a breakthrough by accurately predicting protein structures from amino acid sequences at near-experimental accuracy, solving a decades-old challenge in biology. Its successor, AlphaFold3 [*3*], further improved on this by enhancing the ability to model more complex protein interactions and assemblies. Meanwhile, RFDiffusion [*4*] and Chroma [*5*] introduced diffusion-based generative models that enable the creation of novel protein structures with desired properties and functions, pushing the boundaries of de novo protein design.

However, the rapid development of protein generative models (PGMs) and the lack of corresponding regulations lead to **Biosecurity** and **Intellectual Property (IP) infringement** concerns. **First**, the unregularized yet powerful protein generative models are vulnerable to misuse and cause bio-security/safety concerns [*7, 8*] (Figure. 1a). For example, such models can be misused to design proteins with harmful properties, including pathogenic, toxic, or viral agents that could be exploited as bioweapons [*9, 10, 11*]. Recently, researchers at Microsoft discovered the ability of open-source AI protein design models to create variants of proteins of concern that could evade detection by the biosecurity screening tools [*12*]. These concerns have spurred community-wide discussions and policy dialogues [*7, 8, 13, 14*], further amplified by international biosecurity forums such as those organized by IBBIS, NTI, and the U.S. National Academies of Sciences [*15, 16, 17*]. **Second**, the ease of model sharing brings up IP infringement concerns, including the risk of unauthorized use of generated structures and redistribution of pretrained models for profit, which could undermine the interests of the original creators [*18, 19, 20*]. For example, the latest AlphaFold3 Server has the user terms saying that users “*must not use AlphaFold Server or its outputs to train machine learning models*” or “*in connection with any commercial activities*” [*21*] (Figure. 1b). These restrictions highlight the growing tension between open scientific access and proprietary protection in AI-driven protein research. Therefore, a digital-stage watermarking method would provide a practical mechanism for establishing the provenance of AI-generated protein structures, detecting whether a structure originates from a protected model, and attributing potentially unauthorized or concerning outputs to specific users. [*22, 23, 24, 25*].

**Figure 1.**
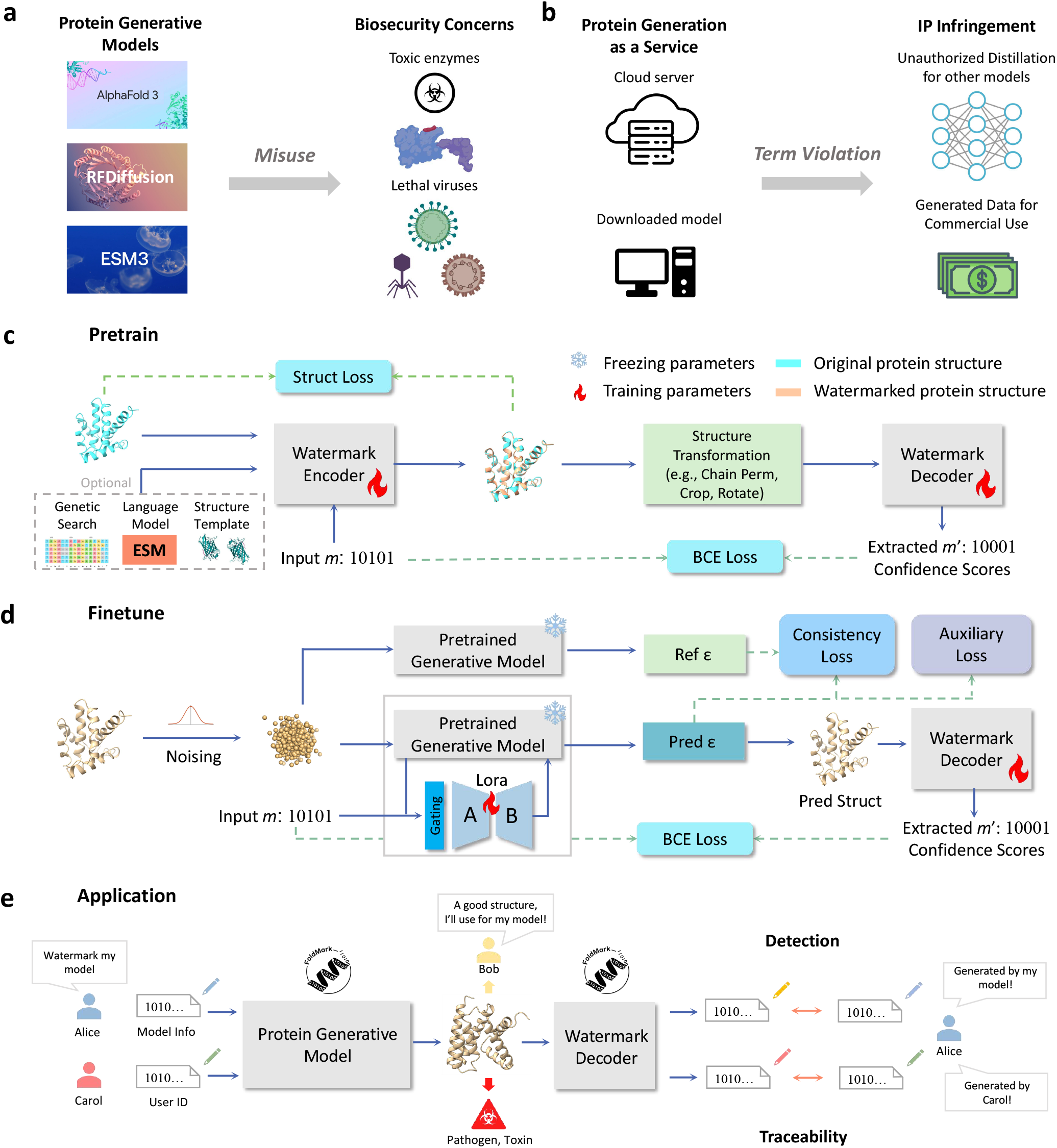
FoldMark: Distributional and Evolutionary Watermark for Protein Generative Models (PGMs). **a**,**b**, Powerful protein generative models, such as AlphaFold3, RFDiffusion, and ESM3, bring increasing biosecurity and IP infringement concerns [*13, 14*]. **a**, PGMs may be misused for synthesizing toxic enzymes or lethal viruses. **b**, Protein generative models like the AF3 Server are evolving into ‘Protein Generation as a Service’ (PGaaS). AF3 Server has terms that users “*must not use AlphaFold Server or its outputs to train machine learning models*” or “*in connection with any commercial activities*” [*21*]. However, users may breach terms via unauthorized distillation for unauthorized commercial use. **c**,**d**, FoldMark provides built-in protection for PGMs by embedding watermarks during protein structure generation for reliable tracking. The implementation of FoldMark on PGMs takes a two-stage process. **c**, FoldMark pretraining: The FoldMark Encoder learns to embed watermark codes by subtly modifying protein structures. It also accepts a wide variety of optional input features, such as genetic search, protein language model embeddings, and structural templates for better performance. FoldMark Decoder learns to recover the watermark based on the watermark structures. **d**, FoldMark finetuning: The pretrained PGM is finetuned with watermark-conditioned LoRA. **e**, Applying FoldMark for generated-structure detection and user tracing. The model owner, Alice releases the pretrained model or deploys it on public platforms. Bob, who downloads Alice’s model and code, generates protein structures and falsely claims ownership. Carol registers as a user on the server and utilizes the API to generate protein structures for pathogen and toxin synthesis. FoldMark allows Alice to embed watermarks in generated protein structures, which can then be extracted to detect and identify unauthorized or illegal use, thus safeguarding biosecurity and intellectual property rights.

Similar problems have occurred in text and image generation. For instance, Large language models (LLMs) such as ChatGPT can be used to create fake news and to cheat on academic writing [*26, 27, 28*]. The latest text-to-image models, such as Stable Diffusion [*29*] and DALL·E 3 [*30*] enable users to create photo-realistic images like deepfakes [*31*] for illegal purposes. As a result, there is growing consensus that the ability to detect, track, and audit the use of AI-generated content is essential for harm reduction and regulation [*27, 32*]. Recently, the watermark has become one of the most promising protection strategies, which embeds hidden patterns in the generated content and is imperceptible to humans, while making the embedded information algorithmically identifiable. Although watermarks have been applied for Language models [*33, 34, 35, 36, 37, 38*] (e.g., SynthID-Text [*37*]) and text-to-image models [*39, 40, 41, 42*] (e.g., AquaLoRA [*41*]), extending these methods to protein structure data presents unique challenges. Unlike text and images, protein structures are highly sensitive to minute changes [*2, 6, 43*], and embedding watermarks without disrupting the biological functionality or stability of the proteins is a complex task. Moreover, protein structures exhibit complex geometrical symmetries, making traditional watermarking methods less effective due to the requirement for equivariance [*4, 44*].

In this paper, as a proof of concept, we propose FoldMark, a generalized watermarking method for protein generative models, e.g., AlphaFold3 [*3*], ESMFold [*45*], RFDiffusion [*4*], and RFDiffusionAA [*46*]. FoldMark builds upon pretrained protein generative models and generally has two training stages (Figure. 1 c and d). In the first stage, FoldMark pretrains a watermark encoder and decoder to subtly modify structures, embedding watermark code across all residues (i.e., *distributional*), guided by *evolutionary* signals (e.g., genetic search, protein language model, structure templates) that minimize noise in conserved regions and exploit flexible regions for enhanced capacity, ensuring retrievability without compromising structural quality. In the second stage, we use watermark-conditioned LoRA [*47*], which flexibly encodes the given watermark code and merges it into the original model weights, without changing or adding extra model architecture. The protein generative model is fine-tuned using three objectives: message retrieval loss, consistency loss, and structure auxiliary loss. The message retrieval loss ensures that watermarks are effectively embedded within the generated structures, enabling the reliable recovery of the embedded codes. The consistency loss promotes minimal impact of watermarking on the quality of the generated structures. Additionally, auxiliary losses, such as pairwise distance losses inherited by AlphaFold3 [*3*] and RFDiffusion [*4*], are incorporated to further enhance structural quality. In practical digital-stage applications, FoldMark enables robust generated-structure detection and user tracing before physical synthesis or experimental validation (Figure 1e). This allows model owners to assert provenance over AI-generated structures and trace the user associated with a potentially pathogenic design at the digital entry point of the design-build-test pipeline, thereby supporting intellectual property protection and biosecurity oversight.

In experiments, we observe that FoldMark achieves nearly **100% bit accuracy**, defined as the proportion of correctly recovered watermark bits from the encoded structure. It also maintains high structural validity, as indicated by scRMSD, scTM, and RMSD metrics. These results demonstrate that FoldMark can reliably embed and retrieve watermarks of up to **32 bits** with minimal structural deviation (Figure 2 and 3). Moreover, FoldMark demonstrates strong robustness against both common post-processing operations (such as rotation, cropping, and coordinate noise) and *adaptive attacks* including *structure purification, model fine-tuning*, and *inverse folding regeneration*. Two fundamental applications are considered: watermark detection and user tracing. In detection, it reliably verifies if a structure originates from a specific model (around **99% TPR @ FPR=**10^−5^). In tracing, it accurately traces the exact user by matching watermarks (up to **1 million users**). Further analysis reveals the correlation between watermark confidence of FoldMark and the structural confidence (e.g., pLDDT), demonstrating that FoldMark adaptively embeds watermarks to balance retrieval accuracy and structural integrity (Figure 4). Finally, we apply FoldMark to the pipeline of **EGFP, Cas13, DHFR, and TEV structure-based redesign** (Figure 5): the redesigned EGFP and Cas13 proteins maintain wildtype-equivalent functionality in wet experiments (avg. 98% fluorescence and avg. 95% editing efficiency retained, respectively) with over 90% watermark detection success rates. We further extend FoldMark to the more stringent enzymatic settings of DHFR and TEV protease, where activity depends on active-site geometry and cofactor or substrate recognition. Notably, **2 of 3 top-ranked watermarked enzyme variants retained function**, as measured by DHFR activity or TEV cleavage efficiency, while maintaining a **100% watermark detection rate**, demonstrating FoldMark’s practical utility for secure and ethical protein engineering.

**Figure 2.**
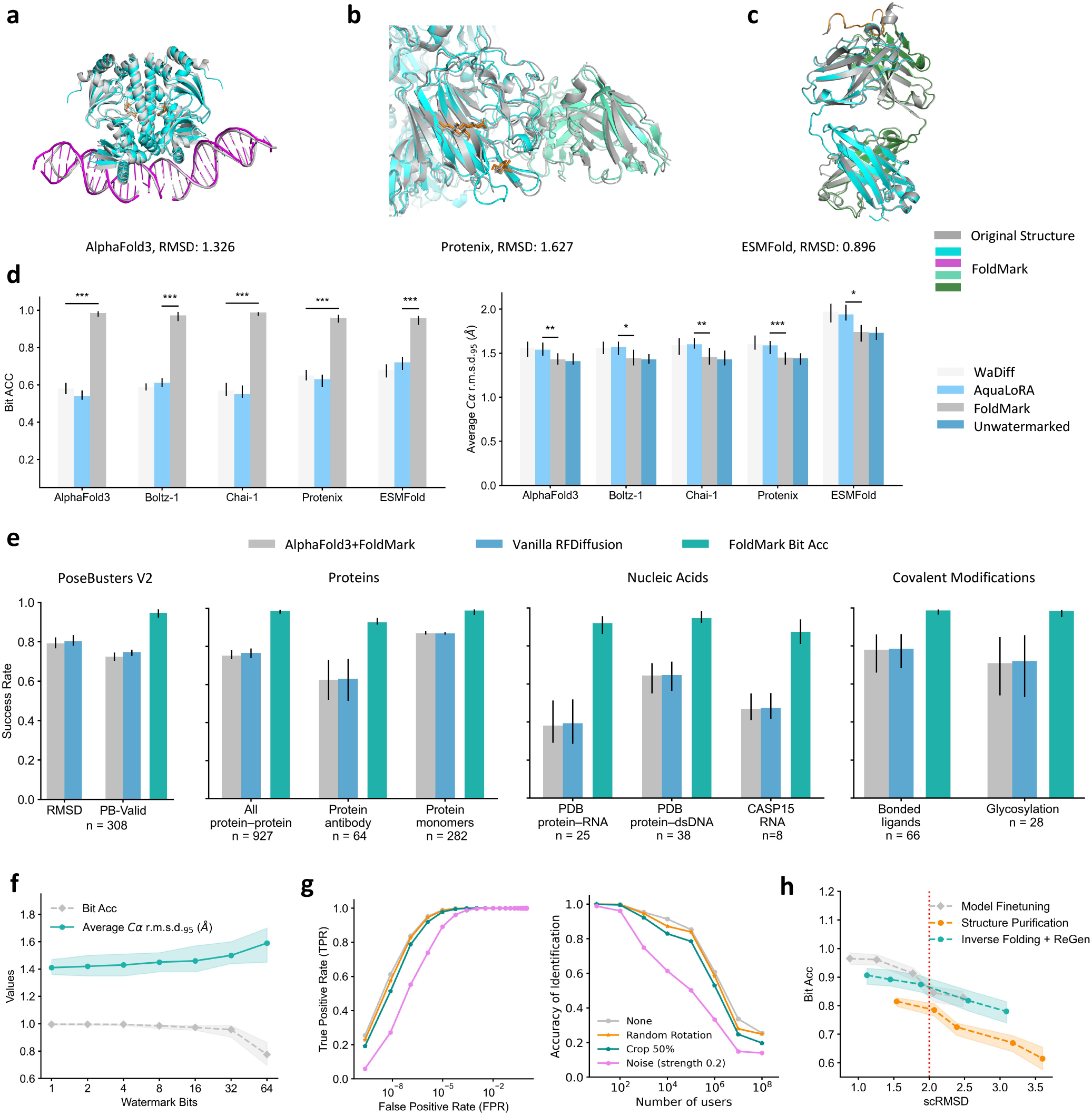
FoldMark Successfully Watermark Structure Prediction Models with Minimal Influence on Structure Quality. **a, b, c**, Example structures watermarked by FoldMark, RMSD is reported between the watermarked and the original predicted structure. **a**, Bacterial CRP/FNR family transcriptional regulator protein bound to DNA and cGMP (PDB 7PZB; Watermarked AlphaFold3). **b**, Human coronavirus OC43 spike protein (PDB 7PNM; Watermarked Protenix). **c**, Mesothelin C-terminal peptide bound to the monoclonal antibody 15B6 (PDB 7U8C; Watermarked ESMFold). **d**, Benchmarking FoldMark with adapted watermark methods from image domain on single protein chains (number of single protein chains = 500). The structural validity is quantified by computing Cα r.m.s.d. at 95% coverage. The means and 95% confidence intervals with 10,000 bootstrap resamplings are shown. Significance levels were calculated using two-sided Wilcoxon signed-rank tests between FoldMark and the second-best watermark method; \*\*\**P* < 0.001, \*\**P* < 0.01, \**P* < 0.05. Exact *P*-values (from left to right) are as follows: 1.46 ×10^−80^, 6.97 ×10^−61^, 1.33× 10^−67^, 2.41× 10^−48^, 3.24 ×10^−20^, 9.73× 10^−3^, 2.11 ×10^−2^, 1.05× 10^−2^, 1.46× 10^−6^, and 4.48× 10^−2^. **e**,The performance of watermarked AF3 with FoldMark on different data categories. Metrics are as follows: percentage of pocket-aligned ligand r.m.s.d. < 2 °A for ligands and covalent modifications; LDDT for nucleic acid and protein monomers; interface LDDT for protein–nucleic acid complexes; and percentage DockQ > 0.23 for protein–protein and protein–antibody interfaces. Sampling and ranking details follow AF3. For ligands, *n* indicates the number of targets; for nucleic acids, *n* indicates the number of structures; for the others, *n* indicates the number of clusters. The bar height indicates the mean; error bars indicate exact binomial distribution 95% confidence intervals for PoseBusters and by 10,000 bootstrap resamples for all others. **f**, AF3 + FoldMark can work well up to 32 watermark bits. The shadows indicate the 95% confidence intervals obtained by 10,000 bootstrap resampling. **g**, FoldMark detection and user identification results with AF3. The left shows TPR/FPR curve of the detection under different transformations. The right is the accuracy of identification with different numbers of users (FPR = 10^−3^). None: no post-processing; Random Rotation: randomly rotated the generated structure; Crop 50%: randomly crop 50% of the continuous protein chains; Noise (strength 0.2): add Gaussian noise with 0.2 strength to coordinates. **h**, Robustness of FoldMark under different adaptive attacks. The red dotted line indicates scRMSD=2.0 and structures with scRMSD under 2.0 are generally regarded as good quality. The shadows indicate the 95% confidence intervals obtained by 10,000 bootstrap resampling.

**Figure 3.**
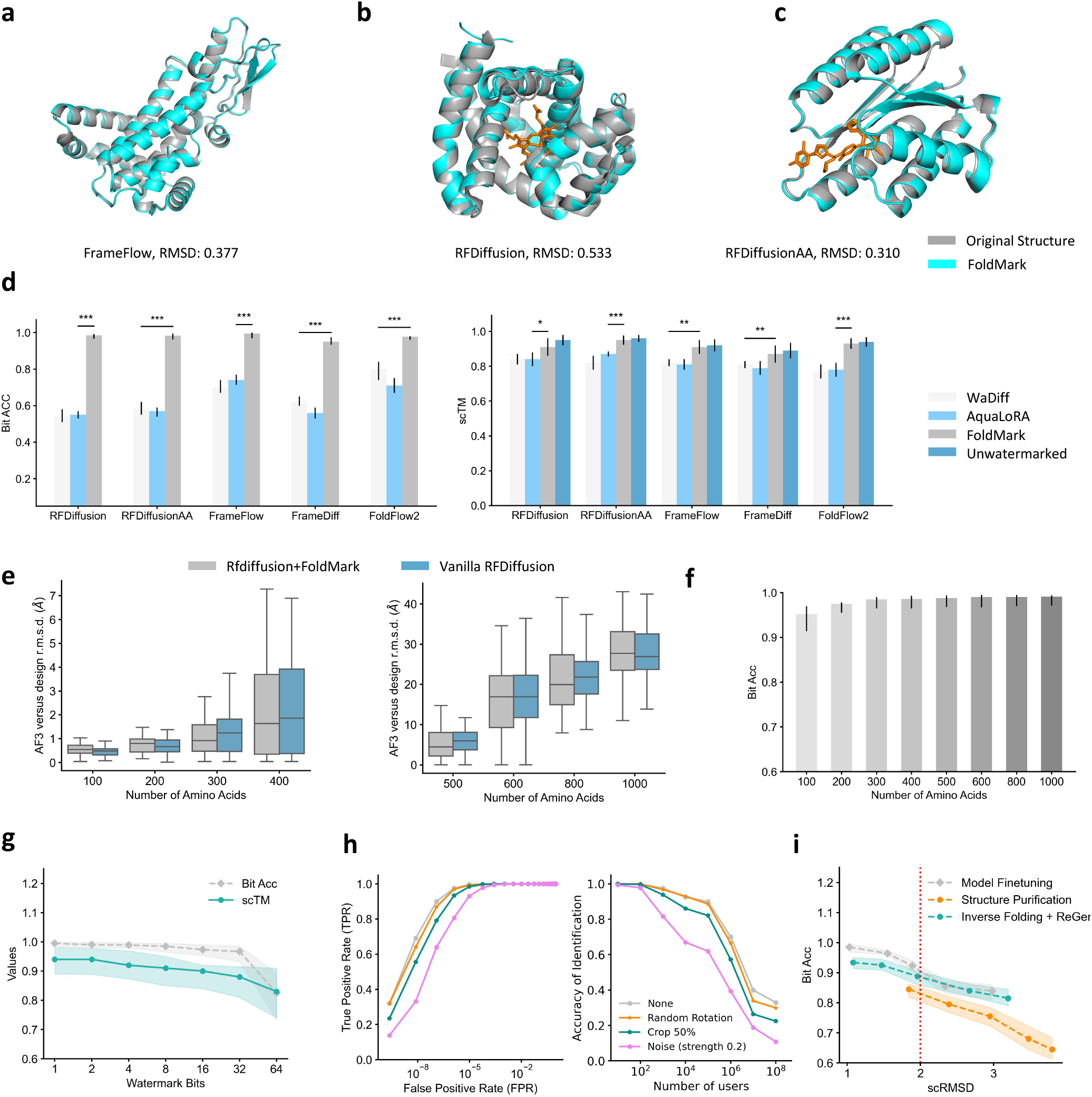
FoldMark Successfully Watermark *de novo* Structure Generation Models. **a, b, c**, Example structures watermarked by FoldMark, RMSD is reported between the watermarked and the original generated structure. **a**, A 256-length protein generated by FrameFlow/ FoldMark+FrameFlow (RMSD:0.377). **b**, Generated binder to the ligand heme from PDB 8vc8 by RFDiffusion/ FoldMark+RFDiffusion (RMSD: 0.533). **c**, Generated binder to the ligand OQO from PDB 7v11 by RFDiffusionAA/ FoldMark+RFDiffusionAA (RMSD:0.310) **d**, Benchmarking FoldMark with adapted watermark methods from image domain on generating 300-length proteins (number of generated proteins = 100). The structural validity is quantified by computing scTM. The means and 95% confidence intervals with 10,000 bootstrap resamplings are shown. Significance levels were calculated using two-sided Wilcoxon signed-rank tests between FoldMark and the second-best watermark method; \*\*\**P* < 0.001, \*\**P* < 0.01, \**P* < 0.05. Exact *P*-values (from left to right) are as follows: 3.89× 10^−18^, 6.91 ×10^−18^, 1.70 ×10^−16^, 3.90× 10^−18^, 1.50× 10^−9^, 3.68 ×10^−2^, 5.55× 10^−7^, 1.09 ×10^−3^, 1.27× 10^−3^, and 8.96 ×10^−8^. **e**, Watermarked RFDiffusion by FoldMark and the vanilla RFDiffusion have similar generation quality (scRMSD) across different lengths. 100 structures are generated at each length. Each box represents the interquartile range (IQR), with the bottom edge marking the 25th percentile (Q1) and the top edge indicating the 75th percentile (Q3). The median (50th percentile) is shown as a horizontal line within each box. Whiskers extend to the minimum and maximum values within 1.5 times the IQR from Q1 and Q3, respectively. **f**, The relationship between watermark capability and the number of amino acids of the generated structure. **g**, RFDiffusion + FoldMark can work well up to 32 watermark bits. The shadows indicate the 95% confidence intervals obtained by 10,000 bootstrap resampling. **h**, FoldMark detection and user identification results with RFDiffusion. The left shows TPR/FPR curve of the detection under different transformations. The right is the accuracy of identification with different numbers of users (FPR=10^−3^). **i**, Robustness of FoldMark under different adaptive attacks. The red dotted line indicates scRMSD=2.0 and structures with scRMSD under 2.0 are generally regarded as good quality. The shadows indicate the 95% confidence intervals obtained by 10,000 bootstrap resampling.

**Figure 4.**
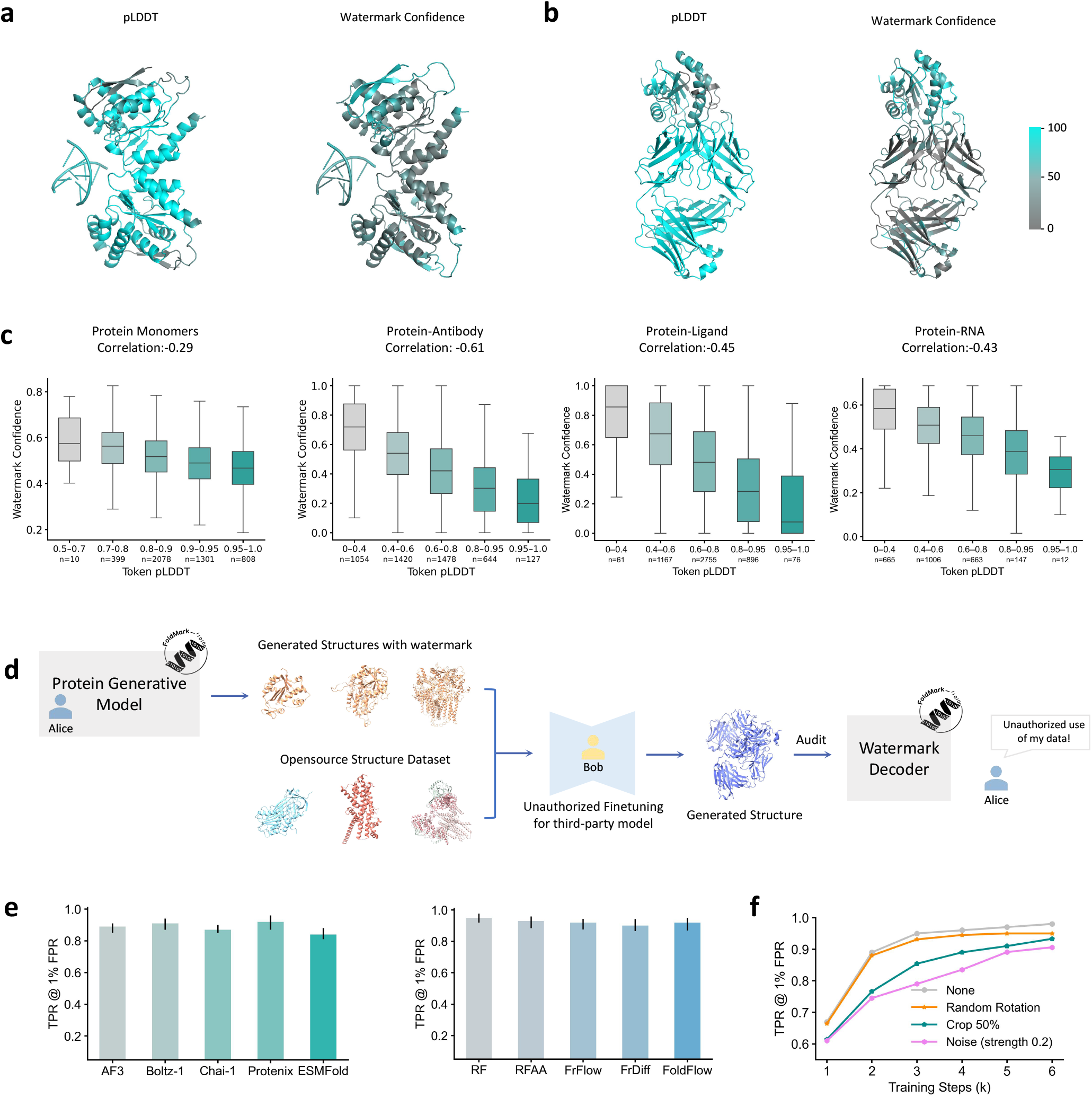
FoldMark’s Distributional Watermark Enables Reliable Detection of Unauthorized Data Use. **a**,**b**, Relationship of Watermark Confidence of FoldMark with Structure Confidence (pLDDT). **a**, Predicted Tir-dsDNA complex (PDB 7×5M; Watermarked AF3). **b**, Predicted structure of HIV-1 MPER scaffold in complex with antibody Fab Ab45.2 (PDB 9BDI; Watermarked Protenix). **c**, The relationship of watermark confidence with token pLDDT. Protein monomers, protein-antibody interfaces, protein-ligand interfaces, and protein-RNA interfaces are selected for evaluation. 50 structures from the first three categories and 25 structures from protein-RNA are selected (100 tokens are randomly sampled in each structure). *n* values report the number of evaluated tokens in each band. The Pearson Correlations are −0.29, −0.61, −0.45, and −0.43 respectively. **d**, FoldMark audits third-party models by detecting inherited watermarks. If a model is trained using protein structures from a watermarked source, its own outputs will contain the original watermark, which FoldMark can recover to identify the unauthorized data use. **e**, Detection True Positive Rate (TPR) across different models at a False Positive Rate (FPR) of just 1%. Each model generates 1,000 samples independently, with the mean and 95% confidence intervals reported. RF: RFDiffusion, RFAA: RFDiffusionAA, FrFlow: FrameFlow, FrDiff: FrameDiff. **f**, TPR @ 1% FPR as a function of training steps in third-party models under different data transformation conditions. None: no post-processing; Random Rotation: randomly rotated the generated structure; Crop 50%: randomly crop 50% of the continuous protein chains; Noise (strength 0.2): add Gaussian noise with 0.2 strength to coordinates. AF3 is used for evaluation.

**Figure 5.**
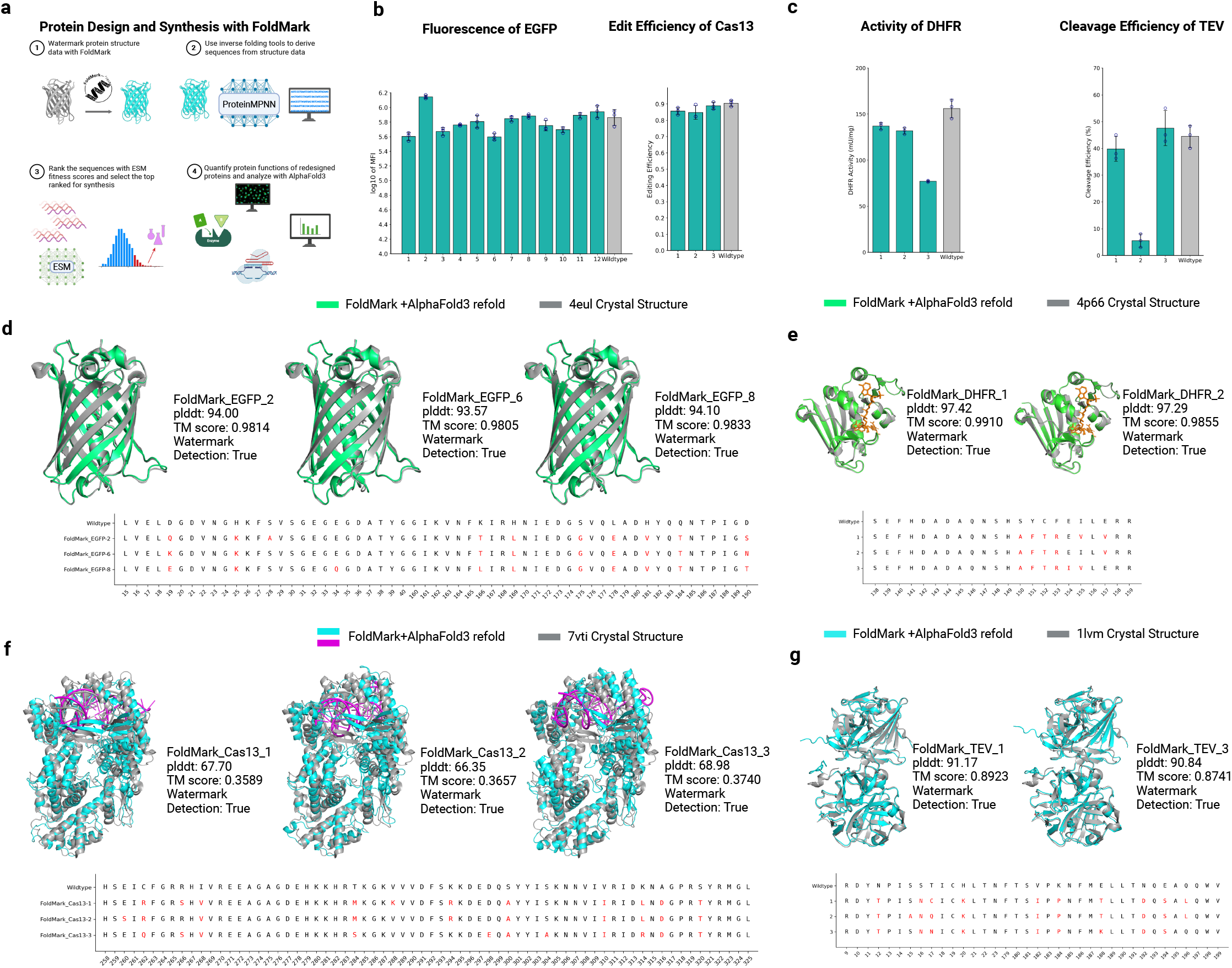
EGFP, Cas13, DHFR, and TEV redesign with FoldMark. **a**, FoldMark-based protein design and synthesis pipeline. Original protein structures are watermarked, redesigned sequences are generated using inverse folding tools, ranked by ESM fitness scores, synthesized, experimentally tested, and analyzed by AlphaFold3-based structural refolding. **b**, Functional validation of FoldMark-redesigned EGFP and Cas13 variants. Redesigned EGFP variants retained an average of 98% fluorescence intensity, and redesigned Cas13 variants maintained an average of 95% editing efficiency relative to wildtype controls. Equivalence to wildtype performance was assessed using two one-sided tests (TOST), with the equivalence margin set to *K* = 3 times the wildtype replicate standard deviation; 7 of 12 EGFP variants and all 3 Cas13 variants met the equivalence criterion (*P* < 0.05). **c**, Functional validation of the top 3 ESM-ranked FoldMark-redesigned DHFR and TEV variants, quantified by colorimetric DHFR activity assay and TEV substrate-cleavage efficiency assay, respectively. **d–g**, AlphaFold3-predicted structures of representative redesigned EGFP, DHFR, Cas13, and TEV variants aligned with the corresponding crystal structures 4eul, 4p66, 7vti, and 1lvm, respectively. pLDDT and TM scores are shown, embedded watermarks were successfully detected, and mutation sites are highlighted in red.

In summary, by integrating distributional and evolutionary design principles for protein structures with advanced generative AI techniques, FoldMark provides an innovative framework for watermarking protein generative models and their outputs. As a digital-stage provenance and attribution mechanism, FoldMark enables traceability and auditing of AI-generated proteins, supporting biosecurity and intellectual property protection.

## Results

### FoldMark Watermarks Protein Structures with Distributional and Evolutionary Information

The rapid advancement of generative AI for protein design raises critical ethical challenges, including biosecurity risks of creating pathogenic proteins (Figure. 1a) and IP infringement concerns due to unauthorized use of generated structures (Figure. 1b), prompting the development of FoldMark as a protein watermarking solution to ensure traceability and IP protection. As a generalized watermarking strategy tailored for protein structure generative models, FoldMark consists of two main stages: Pretraining and Finetuning (Figure. 1c & d).

First, FoldMark pretrains a watermark encoder and decoder to subtly modify protein structures, embedding user-specific watermark code that can be reliably retrieved. FoldMark leverages **evolutionary information** for enhanced watermarking: The FoldMark encoder also supports optional input features like genetic search-derived multiple sequence alignments (MSAs) [*48*], protein language model embeddings from ESM-2 3B [*45*], and structural templates from UniRef100 [*49*] and PDB [*50*] databases. These features capture co-evolutionary signals and species-specific nuances, guiding watermark embedding: *conserved regions receive minimal noise to preserve functional integrity, while intrinsically disordered regions tolerate higher noise*, leveraging evolutionary knowledge for robust and context-aware watermarking. The watermark code **m** is a binary string that is encoded into the structure in a **distributional manner**—across all residues or tokens (with the length of **m** referred to as the number of watermark bits). This approach enhances robustness against cropping and other structural modifications that might otherwise compromise complete watermark retrieval. The watermark decoder take the watermarked structure as input, predicts the watermark code/confidence based on residue/token representations, and take a weighted sum to get the final extracted watermark **m**^′^. In the pretraining, the structure loss and watermark binary-cross entropy (BCE) loss are optimized, encouraging small structure deviations and high watermark retrieval accuracy. To enhance the robustness of watermark decoding, the watermarked structure undergoes structure transformation (e.g., symmetric chain permutation, cropping, and random rotation) before feeding into watermark decoder.

Second, protein generative models are fine-tuned with watermark-conditioned parameter efficient finetuning to preserve generation quality while learning to generate watermarked structures with high recovery rates. As most state-of-the-art protein generative models adopt diffusion-based architecture such as RFDiffusion [*4*] and AlphaFold3 [*3*], Figure. 1 d uses diffusion model fine-tuning as one example, which can be generalized to non-diffusion architectures such as ESMFold [*45*]. During fine-tuning, the watermarked structure is first noised to *t* ~ Uniform [0, 1], initiating a controlled corruption process. The fine-tuned PGM then learns to reverse this noise, iteratively refining the structure by predicting the original watermarked structure from the noised structure. To preserve generation quality and avoid overfitting, a consistency loss is added by comparing the predicted noise (Pred ϵ) and the reference noise from the frozen pretrained PGM (Ref ϵ). Note that a watermark-conditioned LoRA [*47*] is employed during fine-tuning to efficiently embed watermark information into protein structures while reducing computational costs. The approach leverages a low-rank adaptation (LoRA) update, formulated as The approach uses a low-rank update Δ**W** (**m**) = **G** (**m**) ⊙ (**A**× **B**), where the gating vector **G**, derived from a binary watermark code **m** via a linear transformation, modulates the LoRA weight update applied to all linear and attention layers in structural prediction modules of PGM. **A** and **B** are low-rank matrices. This update is added to the original model weights as **W**_watermarked_ = **W** +αΔ**W** (**m**), with α scaling the watermark’s influence, enabling flexible and efficient watermark integration (more details in Methods). Moreover, in finetuning, auxiliary losses such as pairwise distance loss from FrameDiff [*51*] penalizing atomic clashes are considered to further encourage structural quality. Finally, the watermark decoder takes the reconstructed structure based on the predicted noise to predict the watermark code and computes the BCE Loss similar to FoldMark Pretraining.

After fine-tuning, the protein generative model gains the ability to produce structures embedded with a designated watermark while largely preserving the utility of the original model. In a controlled-access deployment setting, such as a server-based prediction or design platform, the released model can encode user ID, request ID, or timestamp information into generated structures to support traceability. For open-source deployment, the watermark-conditioned LoRA weights can be merged into the pretrained weights and only the watermarked model is publicly released.

### Watermarking State-of-the-art Protein Generative Models

In Figure. 2 and 3, we showed experiments of watermarking unconditional protein structure generative models (i.e., RFDiffusion [*4*], RFDiffusion-AA [*46*], FoldFlow [*52*], FrameDiff [*51*], and FrameFlow [*53*]) and protein structure prediction models (i.e., AlphaFold3 [*3*], Protenix [*54*], Boltz-1 [*55*], Chai-1 [*56*], and ESMFold [*45*]). Figure. 2a,b,c and 3a,b,c shows the examples of the generated structures by original PGMs and the watermarked PGMs, demonstrating small structure modifications for watermarking. Furthermore, Figure. 2d and Figure. 3d uses watermark bit accuracy (Bit ACC) to benchmark FoldMark with adapted watermark methods from the image domain (WaDiff and AquaLoRA). With customized geometric data augmentation, powerful structure encoder/decoder, optimized fine-tuning strategy, and evolutionary/template supporting information, FoldMark overperforms baseline methods significantly (p<0.001 with two-sided Wilcoxon signed-rank tests), consistently achieving over 95% Bit ACC across different PGMs. We also quantify the structural validity with C*α* r.m.s.d. at 95% coverage and scTM following AlphaFold2 [*2*] and FrameDiff [*51*]. FoldMark performs better than baseline methods while keeping comparable structural quality. In Extended Data Fig. 1, we further compare FoldMark with sequence-level watermark methods by Chen et al., [*38*] and FoldMark still showed significantly better performance. In Figures 2e and 3e, we stratify the performance of FoldMark combined with AlphaFold3 and RFDiffusion across different biomolecule categories (protein-ligand, protein-protein, nucleic acids, and covalent modifications) or varying sequence lengths (100 to 1,000 amino acids). We observe that FoldMark consistently achieves high predictive accuracy for most interactions and shorter sequences (e.g., < 500 amino acids), while structural quality tends to decline for nucleic acid complexes and longer sequences, due to the increased structural complexity. Note that some underrepresented data categories (e.g., protein-antibody) may exhibit lower performance, representing a limitation that could be addressed in future work through expanded training datasets or refined modeling approaches. In Figures. 2f and 3g, we vary the watermark bit length from 1 to 64 and evaluate its impact on watermark bit accuracy (Bit Acc) and structural validity, measured as *C*_*α*_ root-mean-square deviation (r.m.s.d.) at 95% coverage and self-consistency TM-score (scTM). **In most cases with fewer than or equal to 32 bits, FoldMark achieves nearly 100% bit accuracy in recovering watermark codes from encoded protein structures, with minimal impact on structural validity (**< 1.5 **r.m.s.d. and** > 0.9 **scTM)**. More structural validity evaluations such as Rosetta energy and MolProbity scores are shown in Extended Data Fig. 2. Figure 3e explores watermark capacity as a function of protein sequence length. Using distributional watermark codes, FoldMark demonstrates a steady increase in Bit Acc for longer proteins, attributable to the expanded data space available for watermark embedding. Overall, FoldMark provides a robust and effective framework for protecting protein generative models while preserving structural integrity.

### Robustness against Post-processing and Adaptive Attacks

In real applications, the user may unintentionally use post-processing or intentionally take adaptive attacks to bypass the safeguarding of FoldMark. Here, we consider three common post-processing methods and three adaptive attacks in Figure. 2g & h and Figure. 3h & i. The post-processing methods include (1) **Random Rotation**: randomly rotated the generated structure; **(2) Crop** 50%: randomly crop 50% of the continuous protein chains; (3) **Noise** (strength 0.2): add Gaussian noise with 0.2 strength to coordinates. Adaptive attacks include Structure Purification, Model Finetuning, and Inverse Folding+ReGen. (1) **Structure Purification** is inspired by DiffPure [*57*], which uses a clean-version PGM to first noise the watermarked structure (e.g., 20 steps) and then denoise to obtain the new structure; (2) **Model Finetuning** tries to fine-tune the watermarked model using clean protein data to erase the watermark in PGMs; (3) **Inverse Folding + Regen** means first use inverse folding models such as ProteinMPNN [*58*] to obtain the protein sequences and then use clean models to regenerate the structure based on the derived sequence. Our analysis reveals that FoldMark exhibits robust resistance to both standard transformations and these adaptive attacks. Its resilience to cropping, translation, and rotation stems from its distributional watermark scheme, where watermark information is encoded into each residue rather than localized in specific regions. For instance, cropping only partially reduces the signal without fully erasing it, as the remaining residues retain their encoded information. In adaptive attacks, Model Fine-Tuning and Inverse Folding + Regeneration are less successful. Fine-tuning requires substantial clean data and epochs to overwrite the watermark, yet FoldMark’s pretraining on augmented, watermarked structures embeds the watermark deeply into its parameter space, resisting casual erasure. In Inverse Folding + Regeneration, watermark-associated structural features bias the sequence inferred during inverse folding. We then regenerate structures from these sequences using independent clean structure prediction models (more results in Extended Fig. 3). Even under this standard protein design workflow, the regenerated structures continue to preserve recoverable watermark signals, indicating that the watermark is not merely tied to the original structure file but is retained through the structure-to-sequence-to-structure transformation. Adding noise to a watermarked structure and Structure Purification, can partially disrupt the watermark by introducing randomness that obscures the watermark signal. However, this comes at a steep cost: excessive noise could significantly degrade structural quality, often increasing *C*_α_ r.m.s.d. beyond acceptable thresholds (e.g., >2 °A). This trade-off limits the attack’s practicality, as the resulting structures become biologically implausible. To sum up, FoldMark’s integration of a distributional watermark scheme with strategic data augmentation enhances its robustness to data transformation and adaptive attacks.

### Robust Watermark Detection and User Tracing

As shown in Figure 1e, two fundamental applications of FoldMark: watermark detection and user tracing are considered. Consider a scenario with Alice, the model owner, who trains and deploys a pretrained protein generative model (PGM) with inference code on a platform. Bob, an unauthorized user, downloads Alice’s model to generate protein structures, claiming ownership. Carol, a registered user, generates structures via the platform’s API. In watermark detection, extracting Alice’s watermark from a structure confirms her copyright and flags it as artificially generated. In user tracing, Alice assigns unique watermarks to each user (e.g., Carol), enabling identification by matching extracted watermarks against a database, thus tracing infringement sources. Following previous works [*39, 42*], we formulate these two tasks with rigorous statistical tests:

### FoldMark for Detection

Alice embeds a *k*-bit binary code *m* ∈ {0, 1}^*k*^ into generated structures. The watermark decoder extracts a message *m*^′^ from a structure *x* and compares it to *m*. Detection hinges on the number of matching bits, *M* (*m, m*^′^), with a threshold τ ∈ {0, …, *k*}:

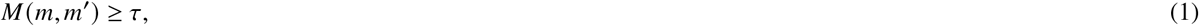

flagging the structure as Alice’s if satisfied. Formally, we test *H*_1_: “Structure was generated by Alice’s model” against *H*_0_: “Structure was not generated by Alice’s model.” Under *H*_0_ (unwatermarked structures), extracted bits 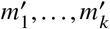 are independent and identically distributed (i.i.d.) Bernoulli random variables with parameter 0.5, so *M* (*m, m*)^′^ follows a binomial distribution with parameters (*k*, 0.5). The True Positive Rate (TPR) is the probability that the watermarked structure is successfully detected; The False Positive Rate (FPR) is the probability that *M* (*m, m*)^′^ exceeds the threshold τ for unwatermarked structures. It is obtained from the cumulative distribution function (CDF) of the binomial distribution and can be expressed using the regularized incomplete beta function *I*_*x*_ (*a*; *b*):

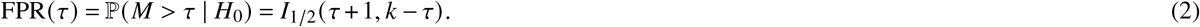

In experiments, Equation 2 is used inversely: we find the threshold τ to achieve a required FPR for detection. We conduct detection experiments with AlphaFold3 and RFDiffusion, generating 5,000 watermarked and unwatermarked structures respectively for each model, and use FoldMark for detection. Figure. 2g and Figure. 3h shows the trade-off between TPR/FPR under different data transformations. We can observe that FoldMark successfully achieves over 90% TPR at FPR equals around 10^−5^, demonstrating robust detection performance even under transformations such as rotation, cropping, and noise addition. More detection results under Amber relaxation [*59*], Rosetta FastRelax [*60*], and NGK loop modeling [*61*] are included in Extended Data Fig. 4.

### FoldMark for Tracing

For tracing, Alice assigns each user *i* (where *i* = 1, …, *N*) a unique *k*-bit signature *m* ^(*i*)^ ∈ {0, 1} ^*k*^, embedded in their model instance. This allows tracking misuse—e.g., identifying a specific user generating viral structures. The extractor compares *m*^′^ from a structure against (*m* ^(1)^, …, *m* ^(*N*)^), testing *N* hypotheses. If all are rejected, the structure is deemed untraceable; otherwise, it’s attributed to argmax_*i*=1..*N*_ *M*(*m*^′^, *m* ^(*i*)^). Regarding the tracing task, false positives are more likely since there are *N* tests. The global False Positive Rate (FPR) at a given threshold τ is:

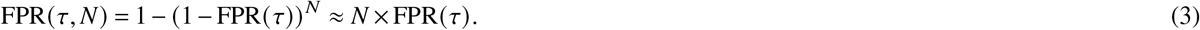

In experiments (Figure 2g and Figure 3h), we vary the number of users and evaluate tracing accuracy, defined as the fraction of watermarked structures correctly attributed to their source user. We conduct these experiments with both AlphaFold3 and RFDiffusion. We perform full simulations without subsampling and scale directly to 10^6^ users [*39, 42*] (see Methods for user-code construction). Each user generates 30 structures, reflecting a more practical usage regime. At a false positive rate of 0.001, FoldMark maintains strong identification performance at large scale, including at the million-user level when structures are not modified. Tracing accuracy remains largely preserved under random rotation and cropping, whereas additive structural noise leads to the largest degradation in tracing performance. Notably, however, such perturbations also substantially impair structural quality. Together, these results demonstrate that FoldMark supports robust and scalable user-level tracing under realistic deployment settings.

### Distributional and Evolutionary Watermark Scheme

FoldMark introduces a novel distributional and evolutionary watermarking scheme, embedding watermark information across all residues of a protein structure, and leveraging evolutionary priors such as MSA and structural to guide the process. In Figure. 4, we conduct a detailed analysis. In Figure. 4a & b, we compare the structural confidence pLDDT from PGMs and the normalized watermark confidence from FoldMark for the predicted Tir-dsDNA complex (PDB 7×5M) and HIV-1 MPER scaffold in complex with antibody Fab Ab45.2 (PDB 9BDI). For 7×5M, the pLDDT is high in the core domains of Tir, such as helical regions involved in membrane anchoring and effector binding, which are evolutionarily conserved and critical regions for function [*62*]. Conversely, the pLDDT is low for flexible loops or linkers connecting Tir’s structured domains and the N- or C-termini of Tir. These regions are often disordered or lack strong evolutionary constraints, leading to low pLDDT. The watermark confidence from Folamark highlights the low-pLDDT regions such as loop regions and assigns low weights to rigid helical regions. Similarly, the pLDDT is high in the conserved MPER helical core and low in antibody CDR loops for 9BDI [*63*]. The watermark confidence has the highest scores for the interface between the Fab and MPER including the CDR loops.

Therefore, FoldMark is an evolutionary information-driven watermarking framework that respects protein context. By leveraging evolutionary signals, it embeds watermarks with precision: conserved regions receive minimal noise to preserve structural integrity, ensuring biological fidelity, while flexible or intrinsically disordered regions tolerate higher noise, increasing watermark capacity and robustness. This adaptive strategy optimizes both detectability and compatibility with protein function. Figure. 4c further explores this across protein monomers, antibody interfaces, ligand interfaces, and RNA interfaces, sampling 100 tokens per structure. The negative Pearson correlations (−0.29, −0.61, −0.45, −0.43) suggest an inverse relationship between watermark confidence and structural confidence (pLDDT), indicating that FoldMark’s distributional watermark subtly modulates less confident, disordered regions more heavily, aligning with evolutionary variability while maintaining detectability.

### FoldMark Detects Unauthorized Model Training with Watermarked Data

With recent advances in protein AI, protein generation as a service (PGaaS) has transformed computational biology. Yet, the availability of such services raises serious IP compliance concerns. For instance, AlphaFold3 prohibits using its data for commercial purposes or for training third-party models, which is a hard to enforce restriction.

FoldMark addresses this challenge and provides a robust audit pipeline enabling the enforcement of commercial usage restrictions, as illustrated in Figure 4d. When the data is misused, for example, if an external party combines watermarked AF3-generated structures with open-source data to fine-tune their own model—FoldMark can recover the embedded watermark, thereby revealing unauthorized use. FoldMark effectively detects watermarks in the outputs of these models, enabling robust auditing. We simulated the auditing process by fine-tuning third-party models with a mixed dataset consisting of 30% watermarked structures and 70% original training data. In Figure. 4e, FoldMark shows robust detection performance across various protein generative models including AF3, Chai-1, RFDiffusionAA, and FrameFlow, achieved high True Positive Rates (TPR) at a False Positive Rate (FPR) of just 1%. Furthermore, Figure. 4f reveals that the TPR steadily increases with additional fine-tuning steps, likely because extended fine-tuning amplifies the watermark signal’s persistence in the model’s output distribution, thereby enhancing FoldMark’s sensitivity.

### Wet-lab Validation of FoldMark in EGFP, Cas13, DHFR, and TEV Structure-based Redesign

To demonstrate the effectiveness of FoldMark in real-world applications, we applied FoldMark to the structure-based redesign of four representative proteins, including **EGFP** [*64*] (PDB ID: 4eul), **Cas13** [*65*] (PDB ID: 7vti), **E. coli dihydrofolate reductase** (DHFR; PDB ID: 4p66), and **tobacco etch virus protease** (TEV; PDB ID: 1lvm), as shown in Fig. 5a. The pipeline begins with watermarking the original protein structures using FoldMark, followed by deriving novel protein sequences from the watermarked structures using inverse folding tools, including ProteinMPNN [*58*]. The designed sequences are then ranked based on ESM fitness scores [*45*], and top-ranked candidates are synthesized for experimental testing.

We first evaluated FoldMark in EGFP and Cas13, two systems with direct functional readouts. For EGFP, FoldMark was applied to loop regions, whereas for Cas13, FoldMark was applied to the lid domain [*66*]. Wet-lab validation demonstrated that the functionality of nearly all redesigned EGFP and Cas13 proteins was comparable to their wild-type counterparts (Fig. 5b). The redesigned EGFP variants retained an average of 98% fluorescence intensity, while the redesigned Cas13 variants maintained an average of 95% editing efficiency, indicating no substantial deviation from wild-type performance. Moreover, two one-sided tests (TOST) for equivalence against wild-type controls, with the equivalence margin set to *K* = 3 times the wild-type replicate standard deviation, indicated wild-type-level performance for **7 of 12** redesigned EGFP variants and for **all 3** redesigned Cas13 variants (*P* < 0.05). Meanwhile, watermark detection success rates exceeded 90%, reaching 91.7% for EGFP and 100% for Cas13.

We next extended FoldMark to enzymatic systems, which represent a more stringent and challenging setting for protein watermarking. Enzymes often require tightly constrained active-site geometries, precise substrate or cofactor recognition, and conformational features that can be sensitive to small sequence perturbations. We therefore selected TEV protease and E. coli DHFR as two complementary enzyme test cases. TEV protease has stringent sequence specificity and thus provides a test of whether watermarking perturbs substrate recognition and catalytic turnover. E. coli DHFR adopts a Rossmann-like fold and depends on coordinated cofactor and substrate binding, making its activity sensitive to subtle conformational changes. For DHFR and TEV, FoldMark was applied preferentially to structurally flexible or surface-exposed regions rather than to the most conserved catalytic motifs. For each enzyme, we synthesized and tested the top-ranked FoldMark variants selected by ESM fitness scores. Functional assays showed that **2 out of 3 top-ranked** variants retained activity equivalent to the corresponding wildtype enzyme under the tested conditions (Fig. 5c). DHFR variants were evaluated using a colorimetric DHFR activity assay, whereas TEV variants were evaluated by measuring cleavage efficiency of an MBP–TEV site–GFP–HA substrate. These results suggest that FoldMark can preserve enzymatic activity in selected high-ranked candidates, while also revealing that enzyme watermarking is more sensitive to sequence and structural perturbations than reporter-protein redesign. More detailed results of Enzyme experiments are shown in Fig. S9.

To further disentangle the contributions of FoldMark’s watermarking capability from the robustness of inverse-folding models, we conducted structural-level evaluations using AlphaFold3 [*3*]. For EGFP, DHFR, and TEV, the redesigned sequences refolded into structures that were largely consistent with their corresponding crystal structures (Fig. 5d,e,g). Redesigned EGFP and DHFR variants showed particularly high structural agreement, with TM-scores close to 1, while redesigned TEV variants retained overall structural similarity to the TEV crystal structure and preserved detectable watermarks. These results indicate that FoldMark-designed sequences can recover the global structural folds of both reporter and enzymatic proteins after inverse folding. For Cas13, the lower TM-scores likely reflect the limitation of AlphaFold3 for CRISPR-associated system prediction rather than a failure of FoldMark-mediated redesign (Fig. 5f). Under the same prediction setting, even the wild-type Cas13 sequence achieved a TM-score of only 0.38, suggesting that the reduced TM-scores of redesigned Cas13 variants are attributable to the difficulty of accurately predicting this CRISPR-associated protein complex. Nevertheless, all redesigned Cas13 variants retained detectable watermarks and maintained wild-type-level editing efficiency in wet-lab assays.

Together, these findings demonstrate that FoldMark can embed verifiable watermarks while preserving protein function and structural fidelity across diverse protein classes, including fluorescent proteins, CRISPR-associated proteins, and enzymes. At the same time, the DHFR and TEV results highlight that enzyme watermarking imposes additional constraints beyond global structural preservation, particularly around active-site geometry, substrate recognition, cofactor binding, and conformational dynamics. We therefore view enzyme watermarking as an important future direction for FoldMark. In future applications, FoldMark-designed candidates can be combined with additional computational screening steps, including fitness prediction, structure prediction, active-site preservation filters, and substrate- or cofactor-binding site analysis, before wet-lab testing. Such integrated screening may further improve the success rate of functional watermarking in highly constrained enzymatic systems.

## Discussion

This paper establishes the feasibility of watermarking protein generative models and their outputs through FoldMark, a novel method leveraging distributional and evolutionary principles. By pretraining a watermark encoder-decoder to embed user-specific data across residues—guided by evolutionary signals that balance noise in conserved and flexible regions—followed by watermark-conditioned fine-tuning, FoldMark preserves structural quality while enabling robust authentication and tracking. Extensive tests on models like AlphaFold3, ESMFold, RFDiffusion, and RFDiffusionAA demonstrate near-100% watermark recovery with minimal impact on structure integrity, even under post-processing and adaptive attacks. FoldMark’s adaptive embedding, aligned with structural confidence (e.g., pLDDT), further bolsters its resilience, offering a powerful solution to ethical challenges such as IP infringement and unauthorized use in protein design.

There are several limitations that we aim to address in future work. First, FoldMark is currently designed primarily for provenance and attribution within digital protein generation workflows. Advanced users may employ protein generative models beyond one step de novo generation, including iterative structure editing, functional optimization, such as improving binding affinity or enzymatic activity, and motif scaffolding to incorporate specific structural elements into larger frameworks. These multistep transformations may progressively modify or obscure the embedded watermark, potentially weakening its detectability after substantial redesign or iterative refinement. Future work could therefore develop adaptive watermarking strategies that explicitly account for downstream editing, optimization, and repeated transformations. Moreover, FoldMark is not currently intended for direct attribution of experimentally determined structures, such as those reconstructed using cryo electron microscopy, X ray crystallography, or NMR spectroscopy, where experimental noise, conformational heterogeneity, and structure refinement procedures may alter or eliminate the original digital watermark signal.

Second, further engineering and validation are needed to support efficient deployment at the scale of millions of users. Although FoldMark is designed to introduce limited overhead during protein structure generation, large scale deployment would require optimized embedding and decoding mechanisms, scalable codebook management, and robust attribution under decoding errors. Potential directions include sparse or quantized watermark representations, distributed decoding infrastructure, and lightweight preliminary authentication procedures, such as hash based screening or partial decoding, followed by full watermark extraction when necessary. Integration with biotechnology platforms and cloud service providers could further facilitate scalable deployment within existing protein generation infrastructures.

Finally, while FoldMark provides a foundation for output level watermarking of protein generative models, it should be understood as one component of a broader provenance framework rather than a standalone solution. In protein generation as a service deployments, cryptographic and metadata based mechanisms, including authenticated APIs, digitally signed records, access logs, and provenance metadata, can provide strong platform level traceability. However, once generated sequences or structures are exported, shared, modified, or incorporated into downstream workflows, such external provenance information may become detached from the artifact. FoldMark addresses this complementary challenge by embedding attribution signals directly into the generated output. Future work will explore unified provenance frameworks that combine platform level cryptographic safeguards with persistent content level watermarking to enable more comprehensive and resilient attribution throughout the protein design lifecycle.

## Methods

Figure. 1 provide an overview of FoldMark. Inspired by previous works [*39, 41, 40*], FoldMark consists of two main stages: Watermark Encoder/Decoder Pretraining and Protein Generative Model Finetuning. The pretraining stage enables the watermark encoder and decoder to learn how to embed watermark information into the structure space and accurately extract it. The finetuning stage equips pretrained protein generative models with watermarking capabilities while preserving their original generative performance (Consistency-preserving). FoldMark is a versatile method that can be applied to various mainstream protein structure generative models. We use a diffusion-based model as an example and present the details of FoldMark below.

### Watermark Encoder/Decoder Pretraining

We first train a watermark encoder *W* and decoder *D* such that *D* can correctly retrieve the watermark message **m** embedded by *W*:

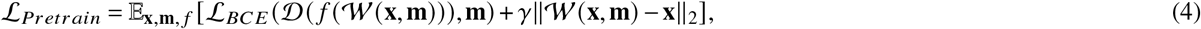

where **x** represents the protein structure data and **m** denotes the string of binary watermark code. γ > 0 is a hyperparameter to control the strength of structure adjustment for watermarking. *f* represents a randomly selected structure transformation as data augmentation. The pool of data augmentation includes random rotation/translation, adding Gaussian noise to protein coordinates, chain permutation, and randomly cropping the protein structure. *L*_*BCE*_ (D (*f* (*W* (**x, m**))), **m** and∥ (**x, m**) −**x**∥_2_ correspond to the CE Loss and Struct Loss in Figure. 1 respectively.

For the watermark encoder/decoder architecture, we have equivariant and non-equivariant versions inspired by prior works [*2, 3*]. The pseudo-algorithms of the water encoder/decoder are shown in the supplementary (Algorithm 1, 2, 3, and 4).

### Optional Evolutionary and Template Input Features

FoldMark Encoder accepts a wide variety of optional input features, such as genetic search, protein language model embeddings, and structural templates for better performance. More details of the use of optional input features are included in the Supplementary.

#### MSA Module

Following Protenix [*54*], we search MSAs using MMseqs2 [*48*] and ColabFold [*67*] MSA pipeline, and use MSAs from the UniRef100 [*49*] database for pairing (species are identified with taxonomy IDs). Retrieved MSA results are cropped to the maximum allowed sequence length. If no hits are found, a length-1 MSA containing only the query sequence is returned.

The MSA module adopted from AF3 processes multiple sequence alignments (MSAs) by embedding a randomly sampled subset of sequences into a representation for each token, updated per recycling iteration. It comprises 4 homogeneous blocks that iteratively refine the pair representation *z*_*ij*_ alongside the MSA, using a network structure similar to the Pairformer Stack, where MSA rows are processed with row-wise attention projected solely from *z*_*i j*_, omitting key-query attention to reduce computational overhead. This design ensures all MSA-derived information flows through the pair representation, enhancing its role as the informational backbone, with updates applied via Triangular Multiplicative Update, Triangular Self-Attention, and SwiGLU-based transition blocks.

#### ESM Module

By training on extensive datasets of protein sequences, protein language models such as Evolutionary Scale Modeling (ESM) are capable of capturing co-evolutionary information inherent in protein sequences. Compared with MSA, ESM eliminates the need for time-consuming sequence retrieval processes and effectively processes sequences with limited homologous data. In FoldMark, we use ESM-2 3B to obtain the per-residue representations for protein chains following Chai-1 [*56*] and add to the initialized single representations.

#### Template Module

Following AF3 [*3*], MSAs derived from UniRef100 are converted into profile hidden Markov models using hmmbuild. These models undergo hmmsearch against a PDB sequence database with stringent parameters (–noali –F 0.1 –E 100 thresholds). Structural eligibility requires minimum length (≥10 residues) and query coverage (>10%), while excluding sequences with >95% identity to the target. Template coordinates are extracted from mmCIF files using sequence alignment validation through Kalign v0.2.0. The system prioritizes templates by e-value ranking, retaining top 20 candidates. During modeling, maximum 4 templates are utilized through randomized selection in training (*k*~ min (Uniform [0, *n*], 4)) versus deterministic top-4 selection in inference.

In FoldMark Encoder, the template embedding integrates all raw template features into a pair representation, which is then processed alongside the existing pair representation *z*_*i j*_. This enables the network to focus on specific regions of the template, guided by its current understanding of the structure.

### Watermark Confidence and Weighted Prediction

In the FoldMark decoder, we extract both a confidence score and a weighted watermark code for robust watermark prediction. The confidence score denoted as *p*_confidence_ = Linear (*a*_*i*_), is computed by applying a linear transformation to the obtained token representations *a*_*i*_. Concurrently, a per-token watermark code *m*_*i*_ = Sigmoid (Linear (*a*_*i*_)) is derived through a separate linear transformation followed by a sigmoid activation, ensuring a bounded output between 0 and 1. The final prediction, *m*_pred_ = Σ_*i*_ Softmax (*p*_confidence_)_*i*_·*m*_*i*_, combines these by weighting each *m*_*i*_ with the corresponding softmax-normalized confidence scores, effectively producing a weighted average that prioritizes contributions based on their predicted reliability. This mechanism enhances the decoder’s ability to integrate watermark information with confidence-driven precision, improving the robustness of the generated outputs.

### Protein Generative Model Finetuning

Instead of finetuning all the parameters of the generative model, we selectively fine-tune part of the protein generative model with LoRA and the watermark decoder as shown in Figure. 1d. The other parameters, such as the reference pretrained model are kept unchanged. We discuss the details of watermark module in the next subsection.

Here, we take the diffusion-based protein generative model (e.g., AlphaFold3 [*3*] and RFDiffusion [*4*]) as an example to construct the fine-tuning loss. The diffusion model typically involves two critical components known as the forward and backward process, where the forward process gradually noises the original protein structure **x**_0_ into **x**_*t*_ for *t* ∈{1, …,*T*}and the model learns to predict the original structure ϵ_θ_ (**x**_*t*_) based on **x**_*t*_. There are two losses in the fine-tuning: the consistency loss for regularization and the message retrieval loss to encourage correct watermark retrieval. In the consistency loss *L*_*c*_, the prediction of the fine-tuned model is compared with the original pretrained model so that the finetuned model weights will not deviate too much from the original ones. For the watermark retrieval loss *L*_*m*_, we take a single reverse step with respect to **x**_*t*_ to obtain 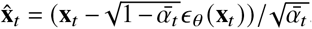, and then feed it into the decoder to predict the watermark code. In sum, we incorporate both optimization objectives above and formulate the consistency-preserving finetuning loss as:

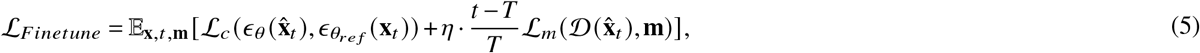

where η controls the trade-off between consistency loss *L*_*c*_ and watermark retrieval loss *L*_*m*_. We place an additional weight 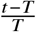 for the retrieval loss because the generated structure contains more information of watermark as *t* →0 and we observe better performance in experiments.

### Watermark-conditioned LoRA

Inspired by previous works in image domains (e.g., AquaLoRA [*41*] and EW-LoRA [*68*]), we use watermark-conditioned LoRA to save the computation costs of fine-tuning and flexibly embed watermark information in the generation process. The computation formula for Watermark-conditioned LoRA in FoldMark can be expressed as:

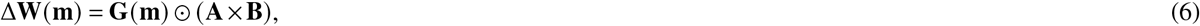

where **A** ∈ℝ^*n*×*r*^ and **B** ∈ℝ^*r*×*m*^ are the low-rank matrices, and **G** ∈ℝ^*n*^ is the gating vector derived from the watermark code. The operator ⊙denotes element-wise multiplication, where **G** modulates the rows of **A**× **B**. This formulation maintains efficiency while allowing flexible incorporation of watermark information.

To input the watermark information into the fine-tuned model, we utilize an adapter layer that converts a watermark code of length *l* into a gating vector **G**. Specifically, the watermark code **m** = {*b*_0_, *b*_1_, …, *b*_*l*_} is passed through a linear transformation defined as:

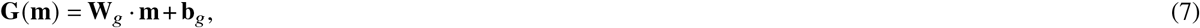

where **W**_*g*_ ∈ℝ^*n*×*l*^ is the linear transformation matrix, and **b**_*g*_ ∈ℝ^*n*^ is the bias vector. Here, *b*_*i*_ ∈ {0, 1} represents the binary state of the *i*-th bit in the watermark code. The gating vector **G** modulates the LoRA weight updates by scaling the rows of the low-rank update **A**×**B**.

During the generation process, when embedding a watermark into the model, we compute the gating vector **G** based on the watermark code. The resulting LoRA weight update Δ**W** is added to the original model weights **W** to produce the watermarked model weights **W**_watermarked_:

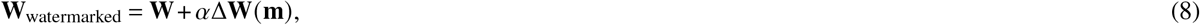

where α is a scaling factor controlling the impact of the watermark on the model weights. FoldMark applies LoRA to all linear and attention layers in structural prediction modules. In contrast, AquaLoRA applies LoRA to linear and convolutional layers in U-Net.

### Training Dataset

#### Dataset for Sturcture Prediction Models

For training of watermarked biomolecule structure prediction models (AF3, Protenix, Boltz-1, and Chai-1), we use all PDB [*50*] structures released before 2021-09-30 (same training cut-off date as AlphaFold3 [*3*]) and with a resolution of at least 9 °Afollowing [*3, 54*]. We parse the Biological Assembly 1 from these structures using their mmCIF file. We use the reference sequence for each polymer chain and align it to the residues available in the structure. For ligands, we use the CCD dictionary to create conformers and match atoms from the structure. We remove atoms under the following conditions: (1) the ligand is covalently bound, and (2) the atom does not appear in the PDB structure. Finally, we follow the same data-cleaning process as AlphaFold3, which includes the ligand exclusion list, the minimum number of resolved residues, and the removal of clashing chains. The total number of PDB structures is around 220k. For the training of watermarked ESMFold, we only use the protein monomer subset.

#### Dataset for Unconditional Structure Generative Models

We trained watermark encoders/decoders and fine-tuned protein generative models using the monomers from the PDB [*50*] dataset, focusing on proteins ranging in length from 60 to 512 residues with a resolution better than 5 °A. This initial dataset consisted of 23,913 proteins. Following previous work [*51*], we refined data by applying an additional filter to include only proteins with high secondary structure content. For each monomer, we used DSSP [*69*] to analyze secondary structures, excluding those with over 50% loops. This filtering process resulted in 20,312 proteins.

### Evaluation Dataset

#### PoseBusters V2

We utilize the PoseBusters Benchmark Set Version 2 (PoseBusters V2) [*70*] to evaluate protein–ligand interactions. This dataset comprises 308 structures. To ensure a fair comparison with AlphaFold3, we adopt their methodology [*3*] to exclude chains that clash with the ligand of interest, specifically in the following PDB entries: 7O1T, 7PUV, 7SCW, 7WJB, 7ZXV, and 8AIE. Similarly, for PDB entry 8F4J, we retain only the protein chains within 20 °A of the ligand.

#### Low Homology Recent PDB Set

We adhere to AlphaFold3 [*3*], Protenix [*54*], and Chai-1 [*56*] to compile the Low Homology Recent PDB set. Homology is defined as sequence identity to sequences in the training set and is measured by template search. Our evaluation set is constructed from a temporal split of PDB entries released between May 1, 2022, and January 12, 2023. All these structures were released after the data in the training sets. We additionally restricted non-NMR structures with a resolution better than 4.5 °A. We restricted to examples that had 2048 or fewer tokens and less than 20 chains. Moreover, we removed monomers and interfaces with substantial homology to the training set:

- Monomers with 40% or greater sequence identity to the training set are removed.
- Polymer-polymer interfaces where both polymers have greater than 40% sequence identity to two chains in the same complex in the training set are filtered out.
- Polymer-peptide interfaces where the non-peptide entity has 40% or greater sequence identity to the training set are removed.

We additionally cluster the above low-homology PDB set. Individual polymer chains were clustered at 40% sequence identity for proteins with more than 9 residues, and 100% sequence identity for proteins with 9 or fewer residues and for all nucleic acids. To study antibody-antigen interface prediction, we filter the low-homology recent PDB set to complexes that contain at least one protein–protein interface where one of the protein chains is in one of the two largest PDB chain clusters (these clusters are representative of antibodies).

We apply quality filters to the bonded ligands and glycosylation dataset (as does PoseBusters): we include only ligands with high-quality experimental data (ranking model fit > 0.5, according to the RCSB structure validation report, that is, X-ray structures with a model quality above the median). As with the PoseBusters set, the bonded ligands and glycosylation datasets are not filtered by homology to the training dataset.

Finally, we obtained 927 protein-protein clusters, 64 protein-antibody clusters, 282 protein monomer clusters, 66 bonded ligand clusters, and 28 Glycosylation clusters. We note that while the above methodology for constructing the low homology recent PDB ret follows the procedure laid out in previous works, we are not able to match exactly the evaluation set used in AF3 [*3*]. As a result, corresponding experimental values on protein-protein interactions, protein-antibody interactions, and protein-RNA interactions may deviate a little from previous works.

#### Filtered Single Chain Set

Based on the clustering of the low-homology PDB set, we select the representative single chain from each cluster, prioritizing chains with a high number of resolved alpha carbons and lengths between 100 and 800 residues. Following AlphaFold2 [*2*], the filtered single chain set is used to benchmark protein structure prediction models in a general manner. The final number of selected protein chains is 500.

#### Nucleic Acids Evaluation Set

For RNA along data, we extracted 8 publicly available targets from CASP15 for evaluation: R1116/8S95, R1117/8FZA, R1126, R1128/8BTZ, R1136/7ZJ4, R1138/[7PTK/7PTL], R1189/7YR7 and R1190/7YR6.

For Protein-Nucleic Acids datasets for evaluation, we screened PDB IDs listed in Table 14 of the AF3 supplementary materials to create the nucleic acid evaluation set. We also omitted systems containing both DNA and RNA simultaneously. From there, we retained only structures with either (i) one RNA chain and one protein chain or (ii) two DNA chains (forming double-stranded DNA) and one protein chain. Next, we filtered out systems with non-standard residues in the DNA. The final dataset matched the counts reported in AF3 Figure 1, consisting of 25 RNA-protein structures and 38 dsDNA-protein structures.

### Metrics

#### Watermark Bit Accuracy

Watermark bit accuracy quantifies the proportion of correctly extracted bits from a watermarked structure, serving as a key metric for evaluating watermarking models. To compute this accuracy, compare the extracted binary bit sequence to the original embedded bit sequence and calculate the ratio of matching bits.

#### scRMSD and scTM

self-consistency *C*_*α*_-RMSD (scRMSD, lower is better) and TM-score (scTM, higher is better) are used to evaluate the quality of generated structures following previous works [*51, 4, 5*]. They measure whether there exists an amino acid sequence that folds to the generated structure and the consistency of sequence-structure pairs. In this work, we use ProteinMPNN [*58*] at the temperature of 0.1 to generate 8 sequences for AF3 [*3*] to predict structures. The final sequence-structure pair is the original sampled structure along with the ProteinMPNN sequence with minimum scRMSD or highest scTM. Generally, a generated structure is called high-quality or designable if scRMSD < 2.0 °A.

#### lDDT

To assess folding accuracy for protein and nucleotide monomers, we employ the Local Difference Delta Test (LDDT) [*71*] computed using specific reference atoms. For proteins, LDDT is calculated on C*α* atoms with an inclusion radius of 15 °A. For nucleotides, it is evaluated on C1^′^ atoms with an inclusion radius of 30 °A. To evaluate protein-nucleotide interfaces, we use the interface LDDT (iLDDT), calculated over C*α* and C1^′^ atoms within the interface.

#### DockQ

We measure the quality of protein-protein interfaces using DockQ [*72*]. If the complex contains multiple identical entities, the optimal assignment of the predicted units to the ground truth units is found by either an exhaustive search over all permutations (for groups up to 8 members) or a simulated annealing optimization (for larger groups) following AlphaFold3 [*3*].

#### Cα r.m.s.d. at 95% coverage

We also report accuracies using the r.m.s.d._95_ (C*α* r.m.s.d. at 95% coverage). To calculate this, we perform five iterations of the following steps: (1) a least-squares alignment of the predicted structure with the PDB structure, based on the currently selected C*α* atoms (using all C*α* atoms in the first iteration), and (2) selection of the 95% of C*α* atoms with the lowest alignment error. The r.m.s.d. of the atoms selected in the final iteration is defined as the r.m.s.d._95_. This metric is more robust against apparent errors that may arise from crystal structure artifacts, though in some cases, the excluded 5% of residues may include genuine modeling errors.

## Implementations

Our FoldMark model is pretrained for 20 epochs and fine-tuned for 10 epochs with Adam [*73*] optimizer, where the learning rate is 0.0001, and the max batch size is 64. We use the batching strategy from FrameDiff [*74*] of combining proteins with the same length into the same batch to remove extraneous padding. In the LoRA, the rank is set as 16 in the default setting. γ and η are set as 2. α is set to 1.0. To minimally influence the generation quality of the pretrained generative models while successfully embedding the watermark, we take an early stop strategy: the training is stopped when the validation watermark bit accuracy is larger than 90%. The evaluation set is constructed by subsampling the training set and the checkpoint frequency is per 500 steps. It takes around 24 hours to finish the whole FoldMark training process (FoldMark pretraining+finetuning) on 1 Tesla H100 80G GPU for AF3. More hyperparameter settings are listed in Table. S4. For each PDB entry, we follow AF3’s inference setup [*3*], generating 25 predictions using 5 model seeds, with each seed producing 5 diffusion samples. The predictions are ranked using confidence scores.

## Watermark Baselines

For comparison, we adapted two state-of-the-art watermarking methods originally developed for image generation: WaDiff [*40*] ^1^ and AquaLoRA [*41*] ^2^. The adapted baselines use their loss functions, training strategy, and part of the watermark model design. Both baseline models were designed for image diffusion models, such as Stable Diffusion [*29*]. Since most protein generative models are also diffusion-based, we applied default hyperparameters from the original works. We also compared with Chen et al., [*38*], a protein sequence watermark methods.

## Implementations of Protein Generative Models

FoldMark is applied for broad state-of-the-art protein structure generative models, including AlphaFold3 [*3*], Protenix [*54*], Boltz-1 [*55*], Chai-1 [*56*], and ESMFold [*45*] for structure prediction, and RFDiffusion [*4*], RFDiffusionAA [*46*], FrameDiff [*51*], FrameFlow [*53*], and FoldFlow2 [*52*] for *de novo* structure generation. We use their open-source pretrained checkpoints and default hyperparameter settings for implementations. More introductions and implementation details are included in the supplementary.

## User-Code Construction

To support reliable attribution at the million-user scale, FoldMark assigns each registered user a unique 32-bit binary signature from a globally planned codebook, rather than independently sampling user codes at random. The codebook is designed to maintain explicit Hamming-distance separation between user signatures, thereby reducing attribution ambiguity when a subset of watermark bits is incorrectly decoded. We first construct a candidate pool using an extended binary BCH-style error-correcting code with parameters [*32, 21, 6*]. This construction yields 2^21^ = 2,097,152 distinct 32-bit candidate codewords and guarantees a minimum pairwise Hamming distance of at least six within the candidate pool. Consequently, any two candidate user signatures differ in at least six bit positions, providing intrinsic tolerance to decoding errors and sufficient capacity for assigning unique signatures to 10^6^ users. From this candidate pool, we construct the final user codebook using a deterministic farthest-first max–min selection procedure. After initializing the selection with a codeword determined by a fixed random seed, we maintain, for each unselected candidate *c*, its minimum Hamming distance to the current selected set *S*, defined as *d*_min_ (*c, S*)= min_*s*∈S_ *d*_H_ (*c, s*). At each iteration, we select the candidate that maximizes *d*_min_ (*c, S*). When multiple candidates have the same minimum distance, ties are resolved by selecting the candidate with the largest average Hamming distance to the existing codebook, followed by a fixed deterministic ordering. This procedure is repeated until 10^6^ user signatures have been selected. Because all selected signatures are drawn from the error-correcting-code candidate pool, the resulting codebook retains a minimum pairwise Hamming distance of at least six, while the max–min selection further improves the empirical nearest-neighbor distance distribution within the million-user subset. Each user identifier is mapped to exactly one codeword through a fixed lookup table, and the complete construction procedure, random seed, and codebook version are retained to ensure reproducible encoding and attribution across model releases.

## Protocols of Wet Experiments

### Structure Templates

Template structures for wet-lab validation were selected to provide high-quality and reproducible structural scaffolds for watermarking and redesign. Selection criteria included experimental resolution, model completeness, local structural definition of the targeted watermarking regions, suitability for downstream experimental validation, and input from wet-lab collaborators with relevant domain expertise. Based on these criteria, we selected **EGFP** [*64*] (PDB ID: 4EUL), **Cas13** [*65*] (PDB ID: 7VTI), ***E. coli* dihydrofolate reductase** (DHFR; PDB ID: 4P66), and **tobacco etch virus protease** (TEV; PDB ID: 1LVM) as templates for the wet-lab validation experiments.

### Watermark Variant Vector Construction

Gene fragments containing watermarked EGFP and truncated Cas13bt3 sequences were synthesized by Twist Biosciences and amplified using Phanta Flash Master Mix. Amplified fragments were gel-purified (NEB) and inserted downstream of the pcDNA3.1 vector backbone by Gibson Assembly using NEBuilder HiFi DNA Assembly Master Mix (NEB). Ligated plasmid products were transformed into competent Stbl3 Chemically Competent *E. coli* (Zymo Research) and validated by Sanger sequencing (Azenta).

His6-TEV and DHFR-His6 genes were synthesized by Twist Biosciences and inserted into PET-28(a) vector bone via NEBuilder HiFi DNA Assembly Master Mix (NEB). The obtained plasmids were validated through whole plasmid sequencing (Plasmidsaurus).

### Mammalian Cell Culture

HEK293T cells (ATCC) were cultured in complete Dulbecco’s Modified Eagle’s Medium (Gibco, 11965-092) supplemented with 10 % (v/v) fetal bovine serum (Gemini Bio, 100-106-500) and 1 % penicillin-streptomycin (Gibco, 15-140-122) at 37 °C under 5 % CO_2_.

### Evaluation of Watermarked EGFP Variants

HEK293T cells were seeded into 96-well plates (Corning) at a density of 2.5× 10^4^ cells per well 24 hours prior to transfection. On the day of transfection, 50 ng of watermarked EGFP plasmids with guide RNA variants were transfected using PEIMAX (Polysciences, 24765). At 24 hours post-transfection, EGFP expression levels were measured by flow cytometry using a CytoFLEX analyzer (Beckman).

### Evaluation of Watermarked Cas13bt3 Variants

HEK293T cells were seeded into 96-well plates (Corning) at a density of 2.5× 10^4^ cells per well 24 hours prior to transfection. On the day of transfection, 10 ng of reporter plasmid (a bicistronic construct consisting of constitutively active mCherry fused with a downstream mutated EGFP harboring a premature stop codon W57* via a self-cleaving T2A peptide), 45 ng of Cas13bt3 guide RNA expressing plasmid (spacer: 5’-GGGTGGGCCAGGGCACGGGCAGCTTGCCGG-3’), and 45 ng of Cas13bt3 watermarked variant expressing plasmids were transfected using PEIMAX (Polysciences, 24765). At 48 hours post-transfection, EGFP and mCherry expression levels were measured by flow cytometry using a CytoFLEX analyzer (Beckman).

### Protein Expression and Purification

Protein expression and purification were performed according to previously described methods [*75, 76*] with minor modifications. Plasmids were transformed into BL21 competent cells (Zymo), and a single colony was cultured in 5 mL TB medium containing 30 µg/mL kanamycin at 37 °C for 12–14 h with shaking at 225 rpm. The starter culture was then subcultured into 500 mL of freshly prepared TB medium at 37 °C until the OD_600_ reached 0.8–1.0. Protein expression was induced by the addition of isopropyl-β-D-thiogalactopyranoside (IPTG; Thermo Fisher) to a final concentration of 0.5 mM at 16 °C for 16 h. Cells were harvested by centrifugation and resuspended in lysis buffer consisting of 50 mM sodium phosphate (pH 8.0), 200 mM NaCl, 10% (v/v) glycerol, 1 mM β-mercaptoethanol, and 25 mM imidazole. The cell suspension was disrupted by sonication on ice, and the clarified lysate was collected by centrifugation at 12,000 ×*g* for 40 min, filtered through a 0.22-µm membrane, and incubated with 1–3 mL of Nuvia™ IMAC Resin (Bio-Rad) at 4 °C. The resin was washed with 5 column volumes of lysis buffer, and bound proteins were eluted with elution buffer consisting of 50 mM sodium phosphate (pH 8.0), 200 mM NaCl, 10% (v/v) glycerol, 1 mM β-mercaptoethanol, and 250 mM imidazole. Appropriate fractions were pooled, concentrated, and buffer-exchanged using an Amicon^®^ Ultra centrifugal filter (Millipore). The purified protein was aliquoted, flash-frozen in liquid nitrogen, and stored at −80 °C.

### Evaluation of Watermarked DHFR Variants

DHFR activity was measured using the Dihydrofolate Reductase Assay Kit (Colorimetric; Abcam, ab239705) with 1 µg of DHFR protein per reaction, following the manufacturer’s protocol at room temperature. To ensure measurement accuracy, all proteins and substrates were used without repeated freeze–thaw cycles. OD_340nm_ was determined using a Tecan plate reader software interface (version 3.9.1.0). The linear regression and *R*^2^ of the NADPH standard curve were calculated using Prism software (version 10.2.2).

### Evaluation of Watermarked TEV Variants

To evaluate TEV protease activity, 10 *µg* of TEV protein was incubated with MBP–TEV site–GFP–HA fusion protein at 4 °C for 16 h in a buffer containing 50 mM Tris-HCl (pH 8.0), 150 mM NaCl, 1 mM DTT, and 0.5 mM EDTA. Reaction mixtures were combined with 4 ×Laemmli sample buffer (Bio-Rad), denatured at 95 °C for 10 min, and resolved on Mini-PROTEAN^®^ TGX™ precast gels (Bio-Rad). Proteins were transferred to PVDF membranes using Trans-Blot Turbo Mini 0.2 *µ*m PVDF transfer packs (Bio-Rad). Membranes were blocked with 5% non-fat dry milk in PBST for 1 h, incubated with an anti-HA tag antibody (Abcam, ab9110; 1:2,000 in PBST) for 1 h, followed by incubation with IRDye^®^ 800CW goat anti-rabbit IgG secondary antibody (LI-COR). Blots were imaged on an Odyssey CLx imaging system (version 1.0.11) using default settings and quantified using Fiji/ImageJ 1.54p (64-bit).

## Data availability

This study’s training and test data are linked at the Github repo (https://github.com/zaixizhang/FoldMark).

## Code availability

The source code of this study is freely available at GitHub (https://github.com/zaixizhang/FoldMark) to allow for replication of the results of this study. The HuggingFace demo website for FoldMark is at https://huggingface.co/spaces/Zaixi/FoldMark.

## Acknowledgements

We thank the authors of WaDiff [*40*] and AquaLoRA [*41*] for discussions, implementations, and feedback.

## Author contributions statement

Z.X.Z., L.C., and M.D.W. designed the research. Z.X.Z. and R.F.J. conducted the computational experiments. G.X.X. and X.T.W. conducted the EGFP and Cas13 experimental validation. J.L. and H.Z. designed and performed the enzyme-related experiments. Z.X.Z., R.F.J., G.X.X., J.L., H.Z., L.C., and M.D.W. analyzed the results. Z.X.Z., R.F.J., M.Z., L.C., and M.D.W. wrote the manuscript. All authors reviewed the manuscript.

## Additional information

**Correspondence and requests for materials** should be addressed to Zaixi Zhang, Huimin Zhao, Le Cong and Mengdi Wang.

**Extended Data Fig. 1.**
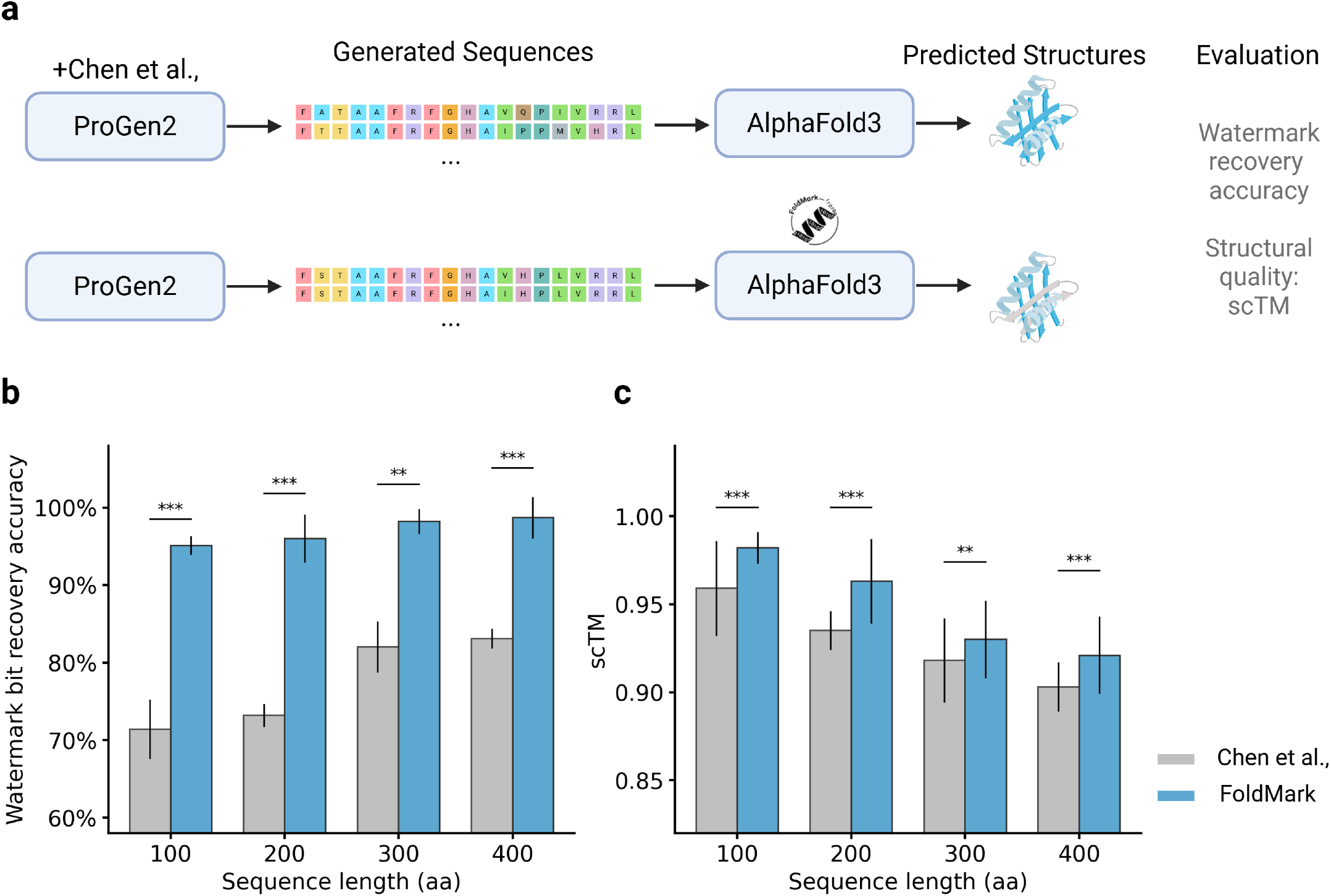
Comparison between FoldMark and the protein-specific baseline of Chen et al., [*38*] under a matched protein generation and evaluation pipeline. Chen et al. implement protein sequence watermarking by modifying the amino-acid sampling distribution during autoregressive protein design according to a private watermark code, so that generated sequences carry a detectable statistical signature while preserving design utility. **a**, Schematic of the comparison setup. Following the setting of Chen et al., protein sequences were first generated using ProGen2 [*77*] under matched conditions, including the same temperature, sequence lengths, and numbers of generated sequences (1000 per length). For the Chen et al. baseline, generated sequences were watermarked at the sequence level and then folded with the original AlphaFold3. For FoldMark, unwatermarked generated sequences were instead folded using AlphaFold3 equipped with FoldMark. The two methods were then evaluated using watermark bit recovery accuracy and structural quality measured by scTM. **b**, Watermark bit recovery accuracy across different sequence lengths. FoldMark consistently achieves higher recovery accuracy than Chen et al. for all tested lengths. **c**, Structural quality, measured by scTM, across different sequence lengths. FoldMark yields higher structural fidelity than Chen et al. across all evaluated settings. Bars indicate mean values and error bars denote s.d. Statistical significance was assessed using two-sided Wilcoxon signed-rank tests. **P < 0.01; ***P < 0.001. These results show that FoldMark consistently outperforms the protein-specific baseline across different protein length regimes, supporting the advantage of structure-stage watermarking over direct sequence-level perturbation.

**Extended Data Fig. 2.**
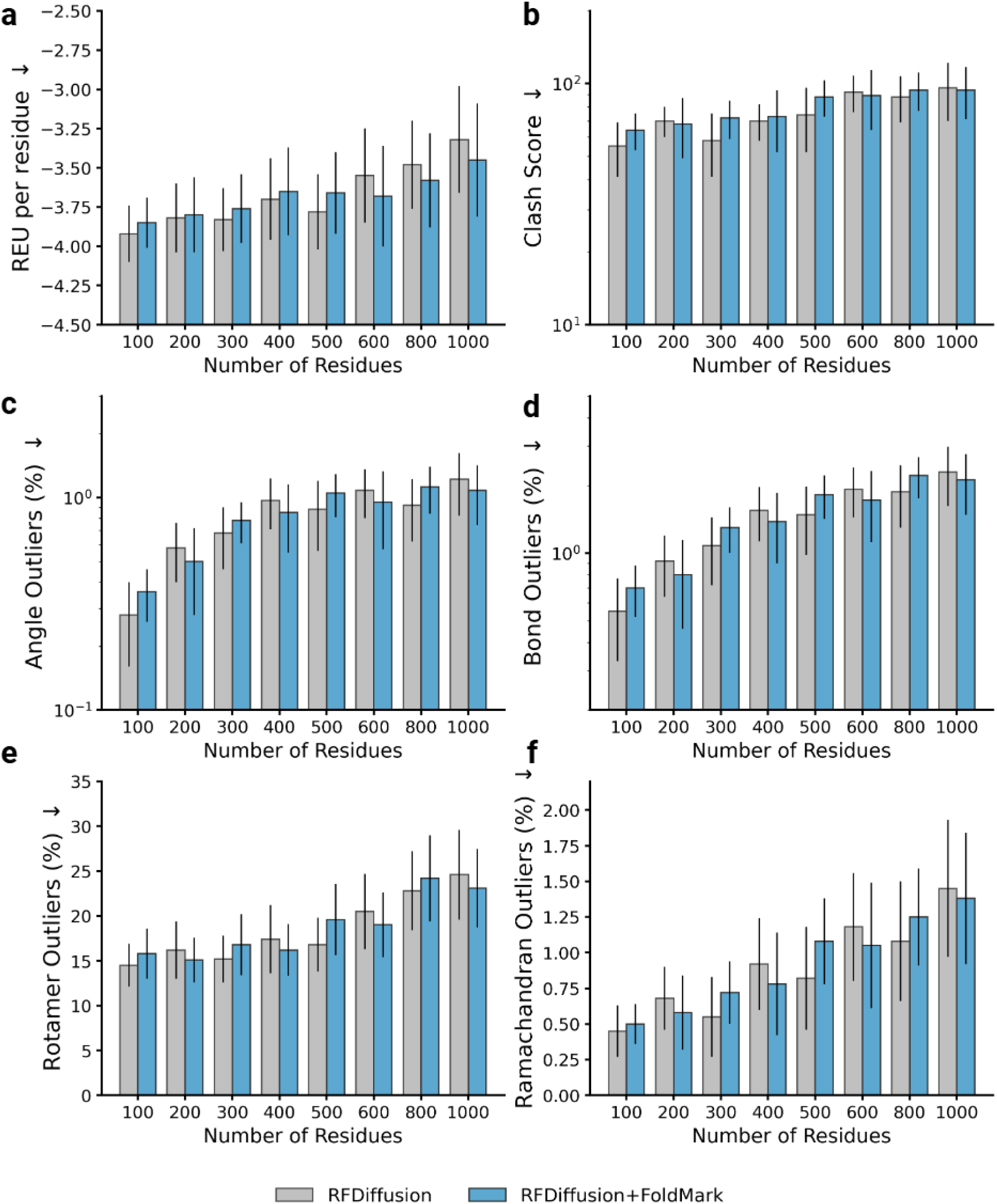
Additional structure validation metrics for RFdiffusion and RFdiffusion+FoldMark across different protein lengths. **a**, Rosetta energy per residue evaluated using PyRosetta 4.0. **b**, Clash score computed using MolProbity. **c**, Angle outliers computed using MolProbity. **d**, Bond outliers computed using MolProbity. **e**, Rotamer outliers computed using MolProbity. **f**, Ramachandran outliers computed using MolProbity. Across all tested protein lengths, RFdiffusion+FoldMark remains highly similar to the original RFdiffusion baseline in all evaluated metrics, indicating that watermarking introduces little to no systematic degradation in structural quality. The results support that FoldMark preserves not only global structural similarity, but also local geometric, stereochemical, and energetic plausibility. Bars indicate mean values and error bars denote s.d.

**Extended Data Fig. 3.**
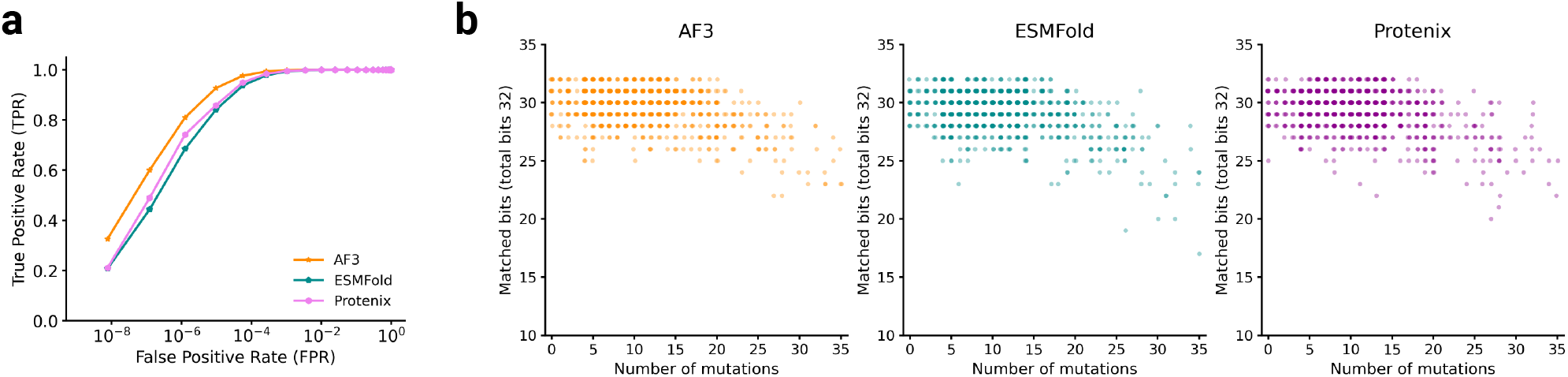
Watermark detection remains robust after inverse folding and clean-model regeneration. **a**, Detection performance of FoldMark after inverse folding followed by clean-model regeneration. Watermarked structures were first converted into protein sequences using inverse folding and then refolded using clean structure prediction models, including AlphaFold3 (AF3), ESMFold, and Protenix. FoldMark maintains high true positive rates (TPR) at low false positive rates (FPR) across all three independent refolding models, indicating that the watermark remains detectable after conversion to sequence and clean-model regeneration. **b**, Per-sample watermark persistence after inverse folding and clean-model regeneration. Each dot represents one inverse-folding-derived sequence. The x-axis shows the number of mutations between the sequence inverse-folded from the original structure and the sequence inverse-folded from the watermarked structure. The y-axis shows the number of matched watermark bits out of a 32-bit binary watermark after refolding with clean structure prediction models, including AF3, ESMFold, and Protenix. Across all three models, sequences with a moderate range of mutations still retain a high number of matched watermark bits, suggesting that the watermark signal can persist through derived sequences and remain recoverable after clean-model refolding.

**Extended Data Fig. 4.**
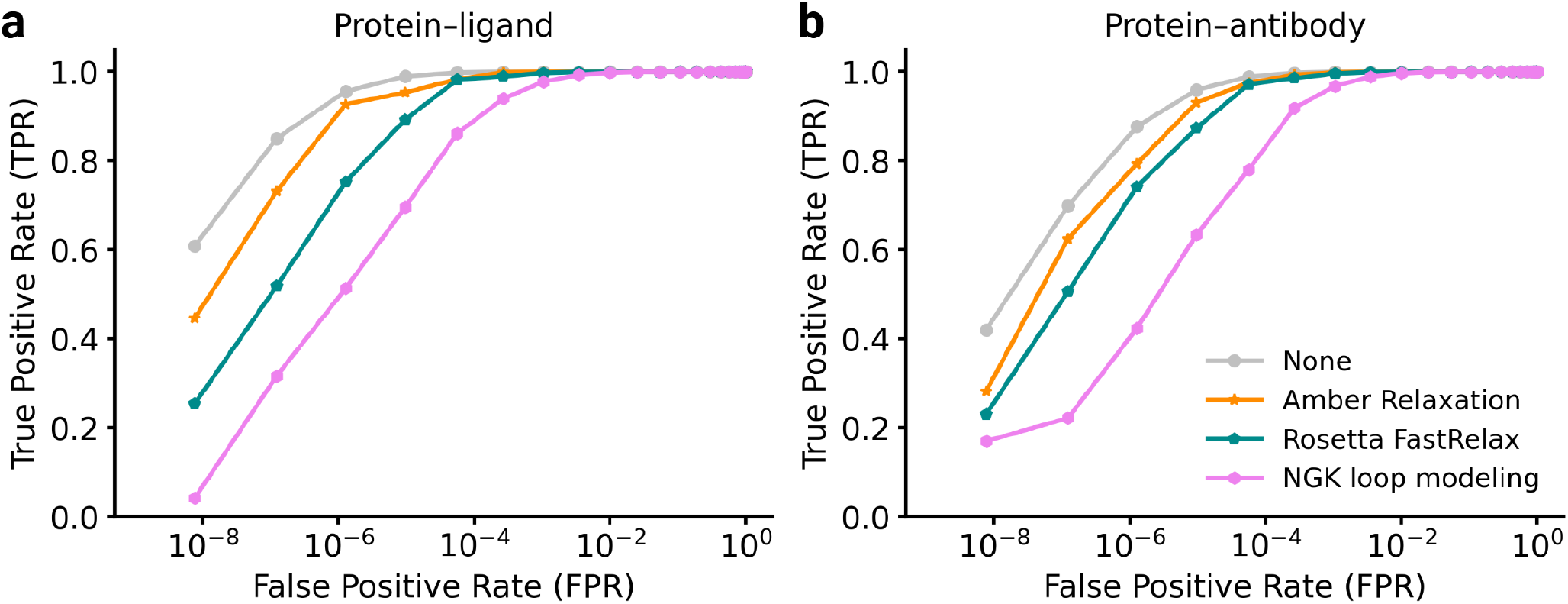
FoldMark remains detectable after structural refinement and local remodeling in bound-state contexts. **a**, True positive rate (TPR) versus false positive rate (FPR) for watermark detection in protein–ligand complexes after Amber relaxation, Rosetta FastRelax, and NGK loop modeling. Unprocessed structures (“None”) are shown as a baseline. **b**, TPR versus FPR for watermark detection in protein–antibody complexes under the same post-processing conditions. In both bound structural contexts, watermark detection is largely preserved after Amber relaxation and Rosetta FastRelax. NGK loop modeling induces a larger decrease in detection performance, as expected from its more aggressive local backbone remodeling, but the watermark remains recoverable overall. These results support that FoldMark is reasonably robust to routine structural refinement and local remodeling in realistic protein complex settings.

**Extended Data Fig. 5.**
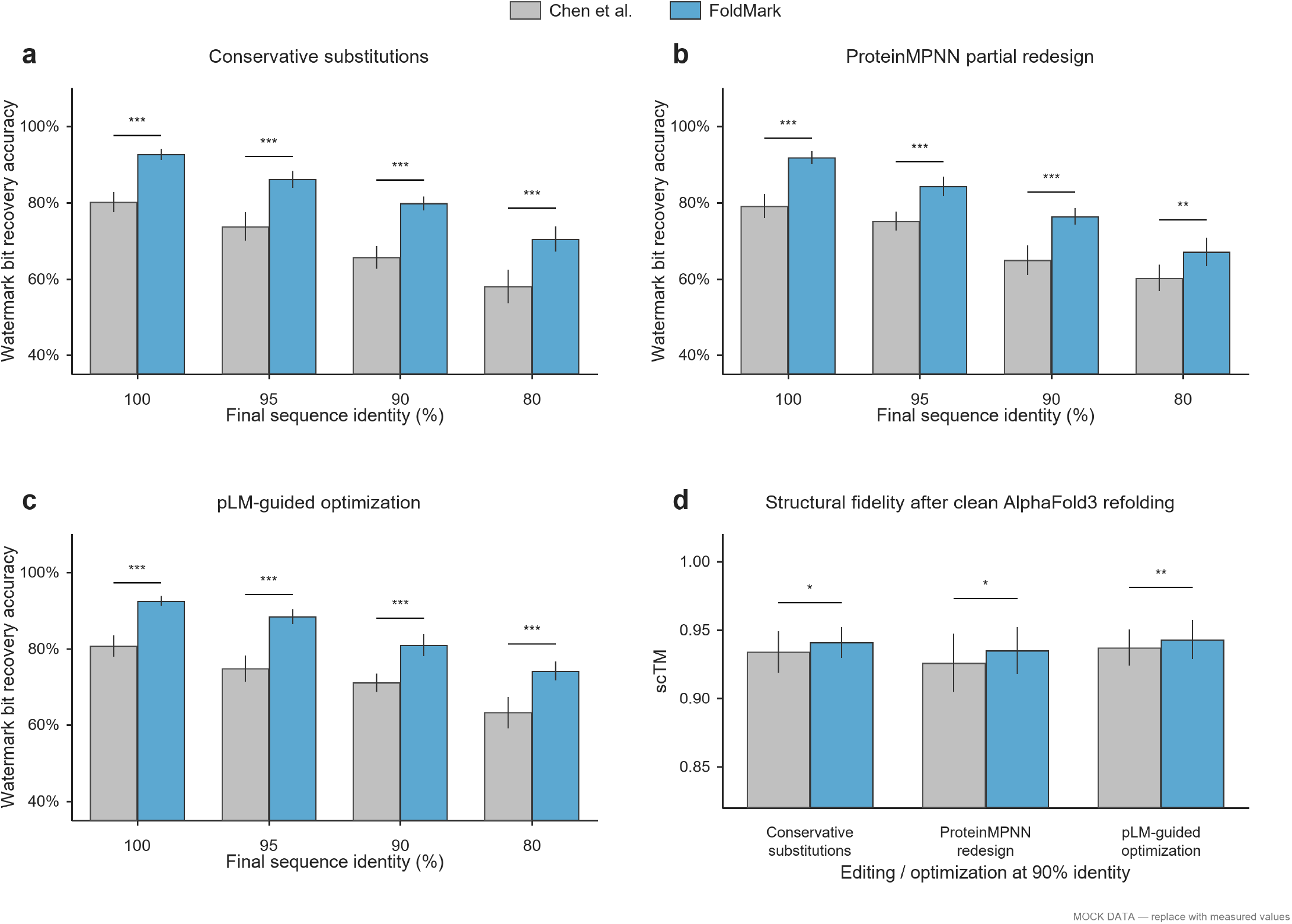
Matched sequence-circulation comparison between FoldMark and the protein-sequence watermarking method of Chen et al. The artifact circulated to downstream users was an amino-acid sequence for both methods. For Chen et al., ProGen2-generated sequences were watermarked directly at the sequence level. For FoldMark, the same source sequences were folded using AlphaFold3 equipped with FoldMark, and the resulting watermarked structures were inverse folded with ProteinMPNN to obtain shareable sequences. Both sets of sequences were subsequently subjected to identical sequence-editing and optimization procedures and then refolded using the original, unwatermarked AlphaFold3 model. **a**, Watermark bit-recovery accuracy after conservative amino-acid substitutions at final sequence identities of 100%, 95%, 90%, and 80%. **b**, Recovery accuracy after ProteinMPNN-based partial sequence redesign at matched final sequence identities. **c**, Recovery accuracy after protein-language-model-guided sequence optimization at matched final sequence identities. **d**, Structural fidelity, measured by scTM, after clean AlphaFold3 refolding at 90% final sequence identity. FoldMark retains significantly higher watermark recovery than Chen et al. across the tested editing operators and sequence-identity levels while maintaining comparable or higher structural fidelity. Bars indicate mean values and error bars denote s.e.m. Statistical significance was assessed using two-sided paired Wilcoxon signed-rank tests: ^∗^*P* < 0.05, ^∗∗^*P* < 0.01, and ^∗∗∗^*P* < 0.001.

## Supplementary Materials

### Materials and Methods

#### Watermark Encoder-Decoder Architecture

##### Algorithm 1

Equivariant FoldMark Encoder

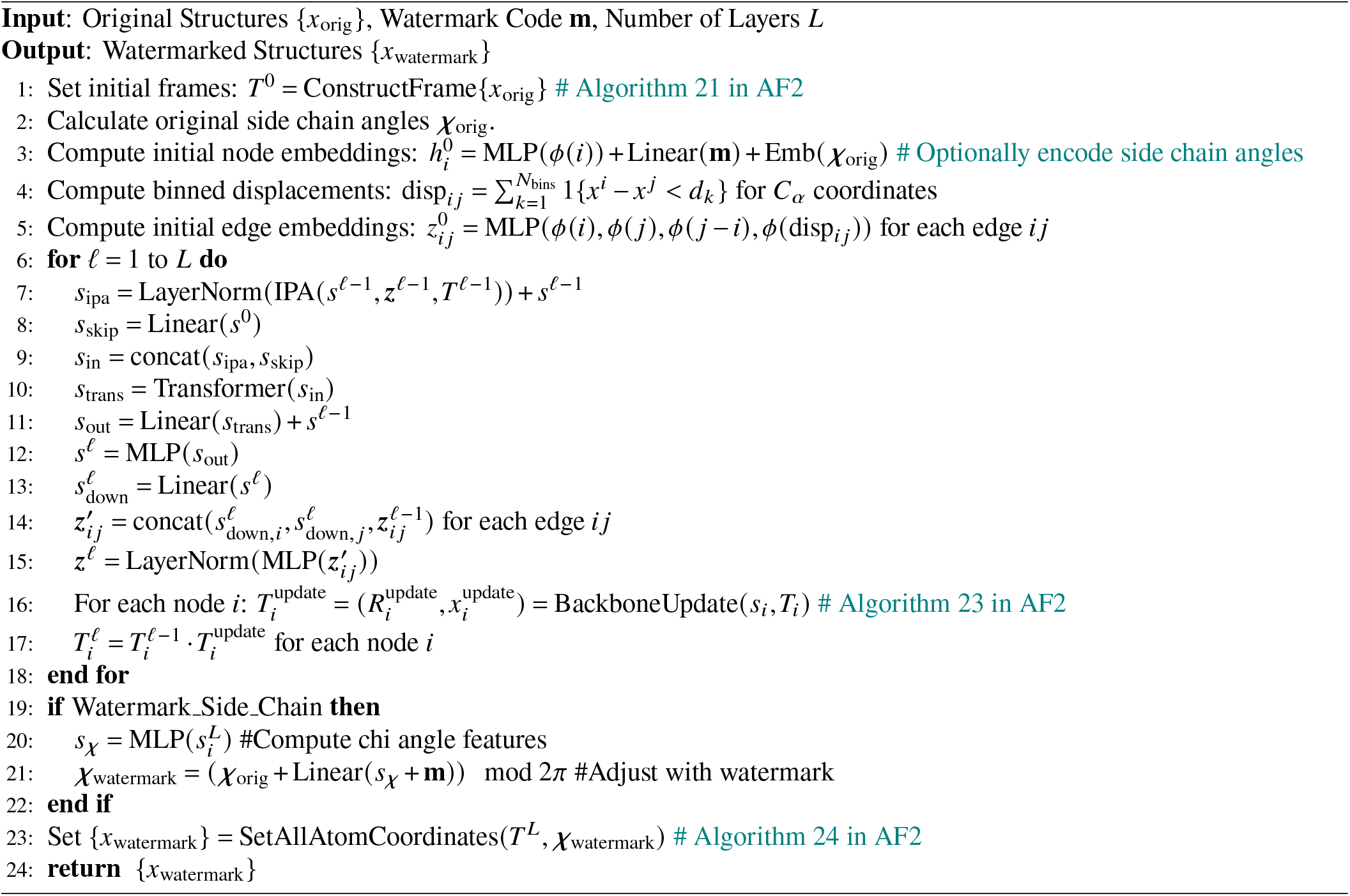

#### Equivariant Architecture

In the FoldMark Equivariant Encoder/Decoder, we adopt a neural network architecture modified from the FrameDiff [*51*]. This architecture consists of Invariant Point Attention from AlphaFold2 [*2*] and transformer blocks [*78*]. Here, for ease of expression, we use superscripts to refer to the network layer and subscripts to indexes or variables. In the complex structure, each residue/ligand atom is represented by one embedding 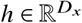 and a frame *T* ∈ *SE* (3). Overall, at the *l*-th layer of the network, 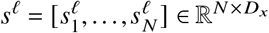 are all the node embeddings where 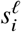 is the embedding for the *i*-th node and *N* is the total number of nodes; 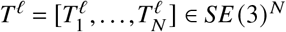 is the frames of every node at the *l*-th layer; 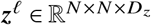 are edge embeddings with 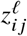 being the embedding of the edge between residues *i* and *j*. In the following paragraphs, we introduce the details of feature initialization, node/edge update, backbone update, and sidechain torsion angle update for watermark encoding and decoding. The pseudo algorithms of Equivariant FoldMark Encoder and Decoder are shown in Algorithm 1 and 2.

#### Feature Initialization

In Equivariant Encoder/Decoder, the node embeddings are first initialized with residue indices while edge embeddings additionally get relative sequence distances. Initial embeddings at layer 0 for residues *i, j* are obtained with an MLP and sinusoidal embeddings ϕ(·) [*78*] over the features. Following [*51*], the binned displacement of two *C*_α_ is given as,

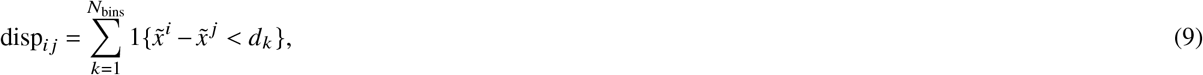

where 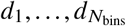 are linspace(0, 20) are equally spaced bins between 0 and 20 angstroms. In our experiments we set *N*_bins_ = 22. The initial node representation is initialized with watermark information:

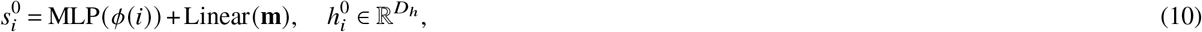

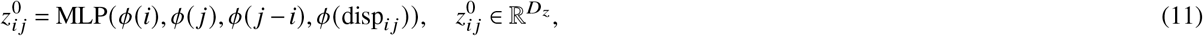

##### Algorithm 2

Equivariant FoldMark Decoder

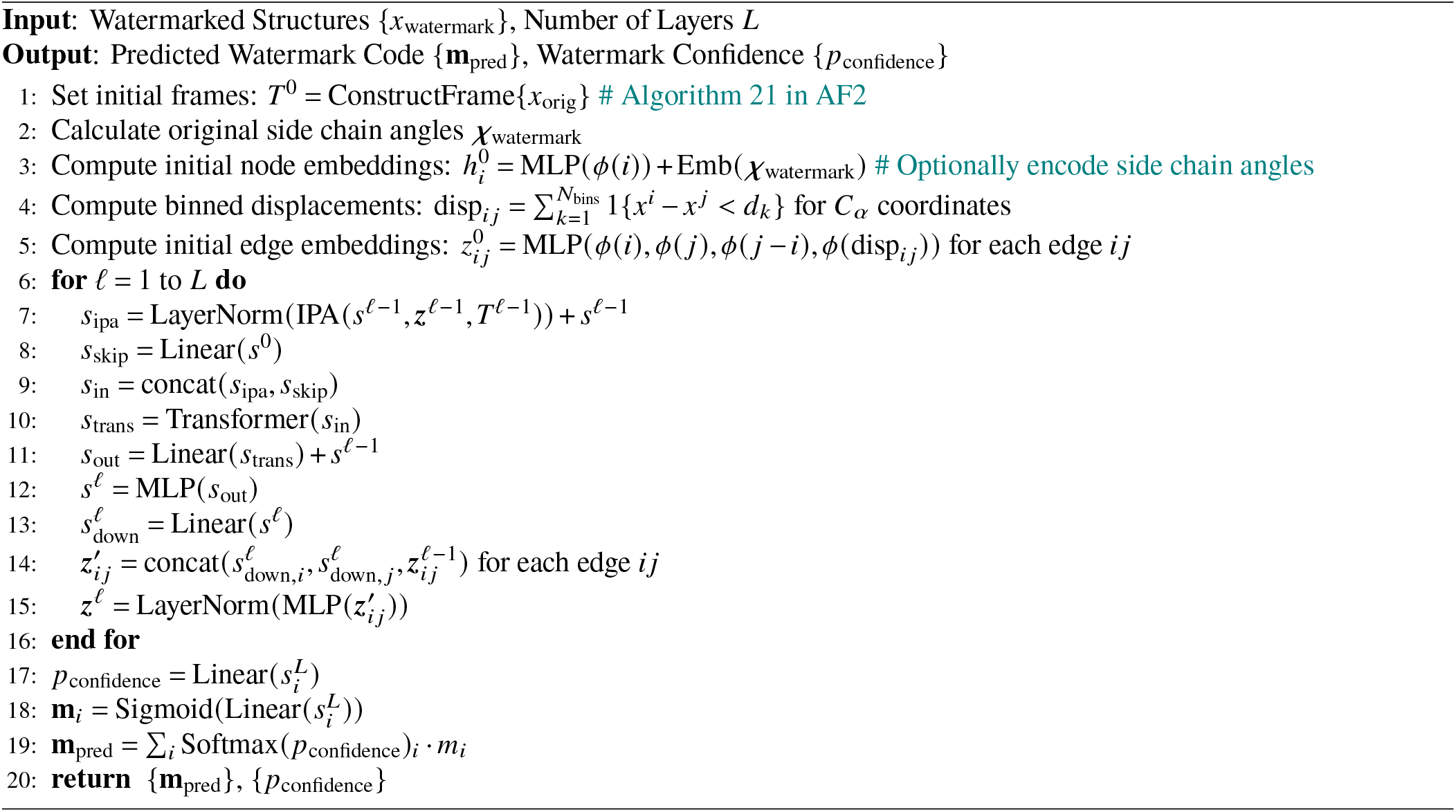

where *D*_*h*_, *D*_*z*_ are node and edge embedding dimensions. If the side chains are considered in the watermarked protein generative models, we may optionally encode side chain angles. We use the input structure to initialize the original frames according to AlphaFold2 [*2*].

#### Node Update

The process of node update is shown below. Invariant Point Attention (IPA) is from AF2 [*2*]. No weight sharing is performed across layers. We use the vanilla Transformer from [*78*]. We use Multi-Layer Perceptrons (MLP) with 3 Linear layers, ReLU activation, and LayerNorm [*79*] after the final layer.

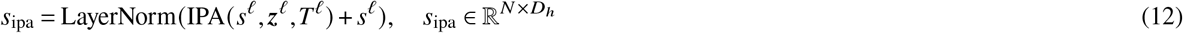

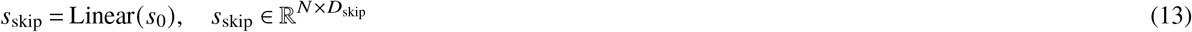

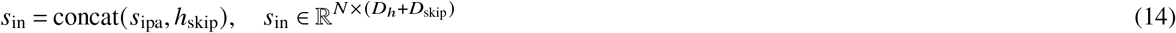

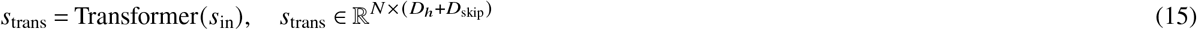

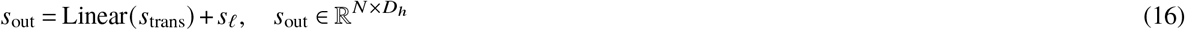

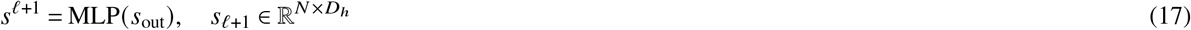

#### Edge Update

Each edge is updated with a MLP over the current edge and source and target node embeddings. In the first line, node embeddings are first projected down to half the dimension.

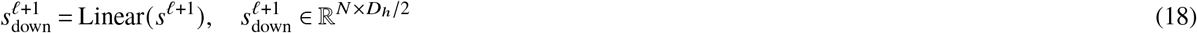

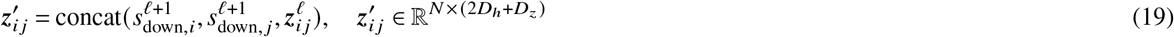

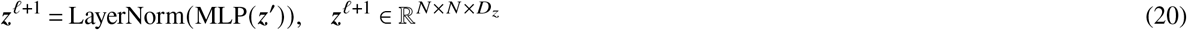

#### Backbone Update

Our frame updates follow the BackboneUpdate algorithm in AlphaFold2 [*2*]. We write the algorithm here with our notation,

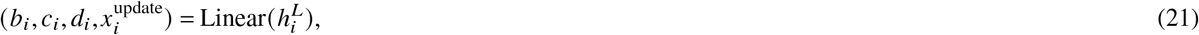

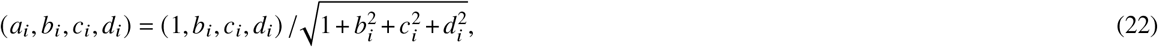

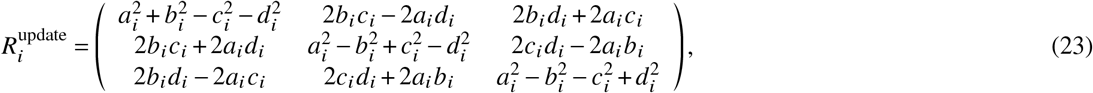

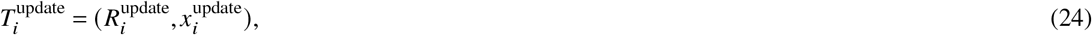

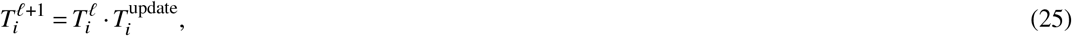

where *b*_*i*_, *c*_*i*_, *d*_*i*_ ∈ ℝ, *x*^update^ ∈ ℝ^3^. Equ. 23 constructs a normalized quaternion which is then converted into a valid rotation matrix in Equ. 24. Following previous works [*53, 4*], we use the planar geometry of the backbone to impute the oxygen atoms.

#### Sidechain Torsion Angle Update

To further encode watermark information, we can update sidechain torsion angles based on watermark code and node embeddings:

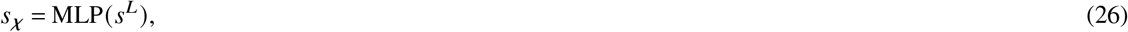

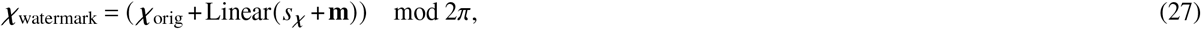

where *χ* ∈ [0, 2π) ^4*N*^. In FoldMark Encoder/Decoder, the number of network blocks is set to 8, the number of transformer layers within each block is set to 4, and the number of hidden channels used in the IPA calculation is set to 16. The node embedding size *D*_*h*_ and the edge embedding size *D*_*z*_ are set as 128. The number of layers *L* for Encoder/Decoder is set to 6. We removed skip connections and psi-angle prediction.

#### Computation of All Atom Coordinates

Following AlphaFold2 [*2*], FoldMark Encoder outputs the watermarked full-atom structure based on the updated backbone frames *T*_*i*_ and side-chain torsion angles *χ* for each residue *i*. These torsion angles are applied to the corresponding amino acid structures, using idealized bond angles and bond lengths, to compute all atom coordinates.

#### Watermark Code and Confidence Prediction

In the FoldMark decoder, we extract both a confidence score and a weighted watermark code for robust watermark prediction. The confidence score denoted as *p*_confidence_ = Linear (*a*_*i*_), is computed by applying a linear transformation to the final node representations 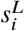. Concurrently, a per-token watermark code 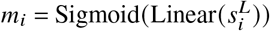 is derived through a separate linear transformation followed by a sigmoid activation, ensuring a bounded output between 0 and 1. The final prediction, *m*_pred_ = Σ_*i*_ Softmax(*p*_confidence_)_*i*_ · *m*_*i*_, combines these by weighting each *m*_*i*_ with the corresponding softmax-normalized confidence scores, effectively producing a weighted average that prioritizes contributions based on their predicted reliability.

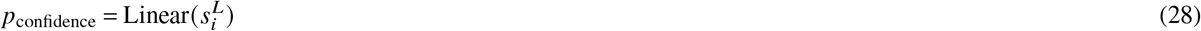

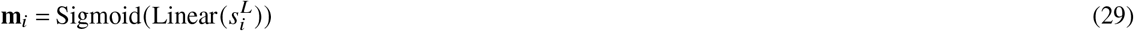

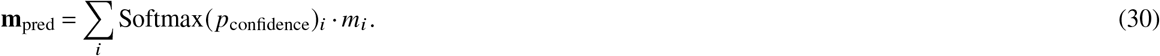

#### Non-Equivariant Architecture

Algorithm 3 and 4 summarize the Non-Equivariant Encoder/Decoder Architecture of FoldMark, which largely follows AF3 [*3*]. The following paragraphs will detail the feature initialization, the feature processing, watermark structure update, and watermark code&confidence prediction.

#### Feature Initialization

For the Non-Equivariant Encoder, the input features follows AF3 [*3*] and can be categorized into six types: Token Features (such as position indexes, chain identifiers, and masks), ESM Features (residue-level embeddings derived from ESM-2 3B), Reference Features (derived from the reference conformer of residues, nucleotides, or ligands), MSA Features (including genetic search-based features like MSA and deletion matrices), Template Features (obtained from template searches for atom positions), and Bond Features (providing bond information, including polymer-ligand and intra/inter-ligand bonds). The reference features use tools like RDKit and fallback to CCD ideal coordinates if necessary. The single and pair features are initialized as:

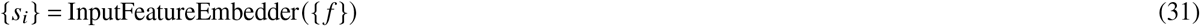

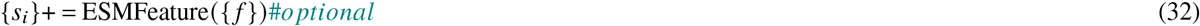

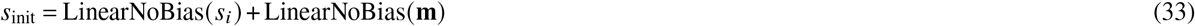

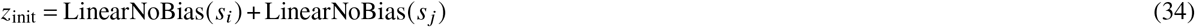

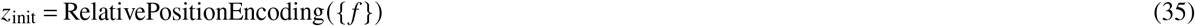

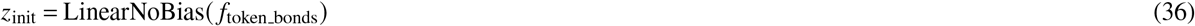

The ESM feature is optional in FoldMark Encoder. For the Decoder, the template, ESM, and MSA input features are removed.

#### Feature Processing

The Encoder mainly contains two parts: the feature processing and watermark structure update. The watermark information is simply encoded with linear layers and added into both modules. In the feature processing part, the single and pair representations are initialized, and encoded information from TemplateEmbedder and MSA Modules. We use MSAs from both protein and RNA sequences. The resulting pair representation is then fed into the main PairformerStack, which also takes the single representation as input. Multiple recycling steps are applied to the aforementioned processing:

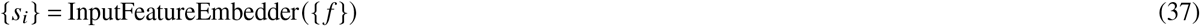

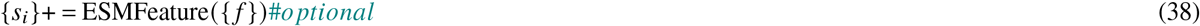

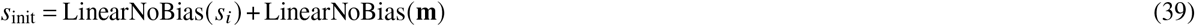

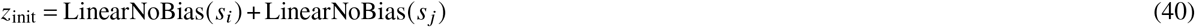

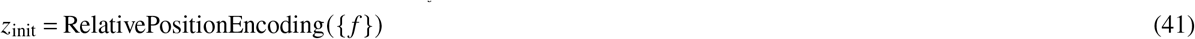

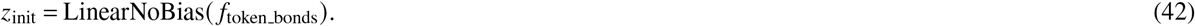

#### Watermark Structure Update

In the watermark structure update, a non-equivariant point-cloud structure prediction module over all atoms from AF3 [*3*] is used instead of IPA in the Equivariant Encoder/Decoder. The structure is first centered and then input to the structure update module to predict the update. The structure module follows a two-level architecture, first processing atoms, then tokens, and finally atoms again. The transformer uses only a single linear layer to embed all atom positions, and a single linear layer to project the watermark updates at the end, without involving any geometric biases (such as locality or SE(3) invariance), therefore non-equivariant. The structure update process is shown below:

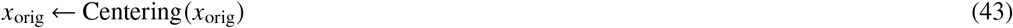

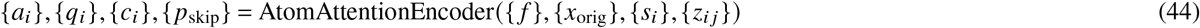

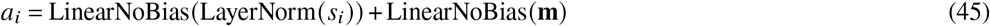

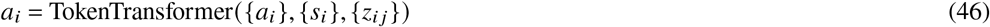

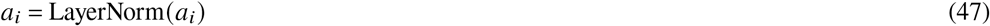

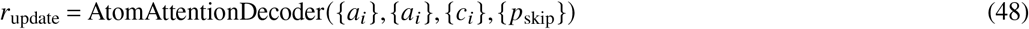

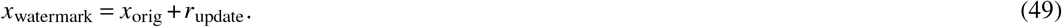

For the decoder, the predicted structure update *r*_update_ is not required.

#### Watermark Code and Confidence Prediction

As for the Decoder, the AtomAttentionDecoder is removed and the per-token watermark code and watermark confidence are first predicted based on the token representations. Then FoldMark

##### Algorithm 3

Non-Equivariant FoldMark Encoder

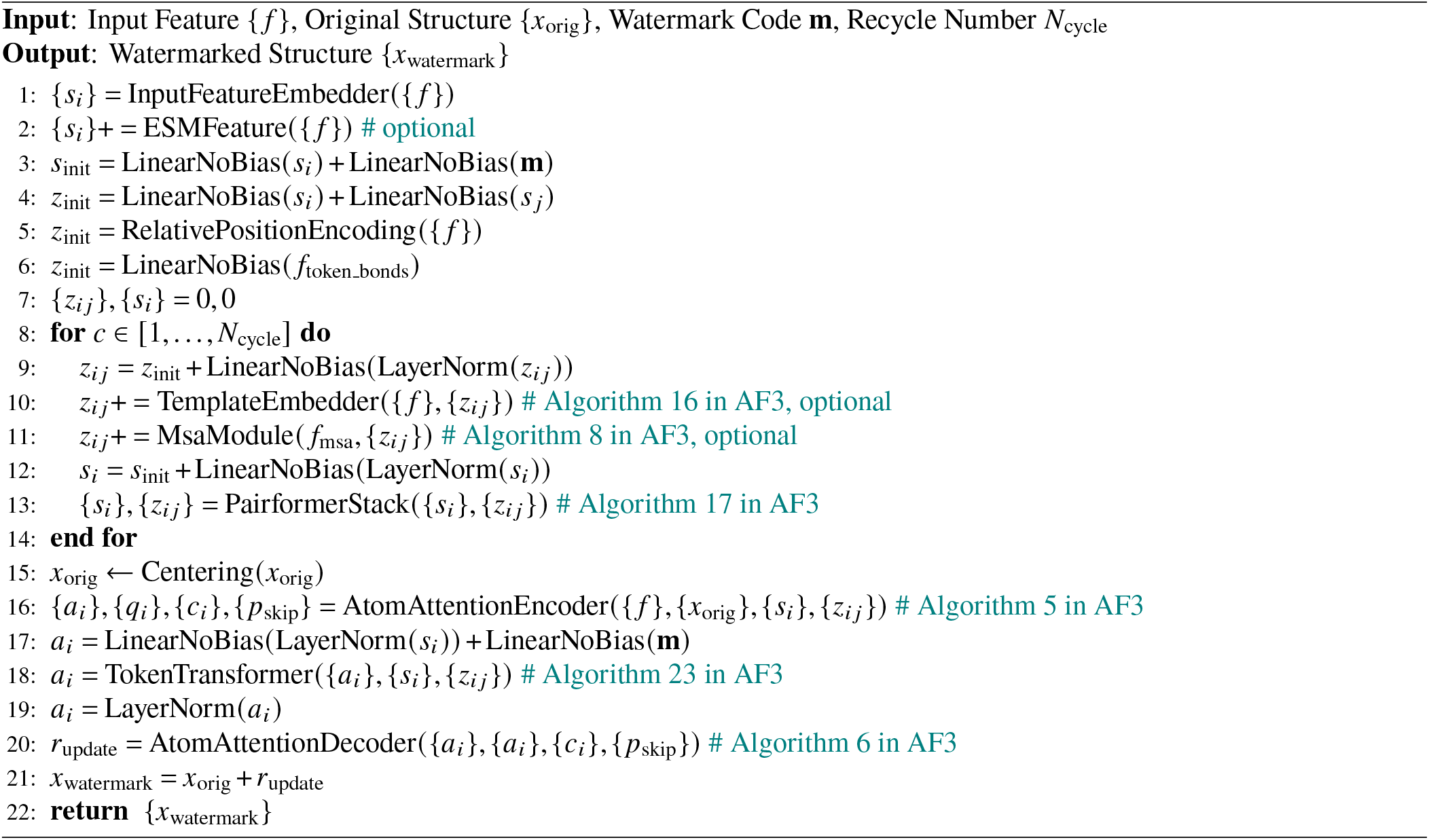

Decoder takes a weighted average to obtain the final watermark code of the whole structure:

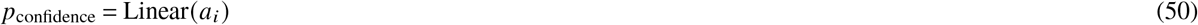

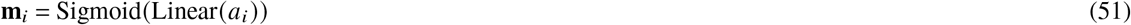

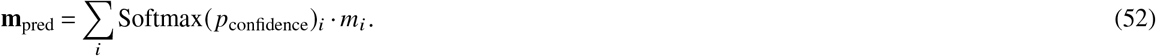

##### Algorithm 4

Non-Equivariant FoldMark Decoder

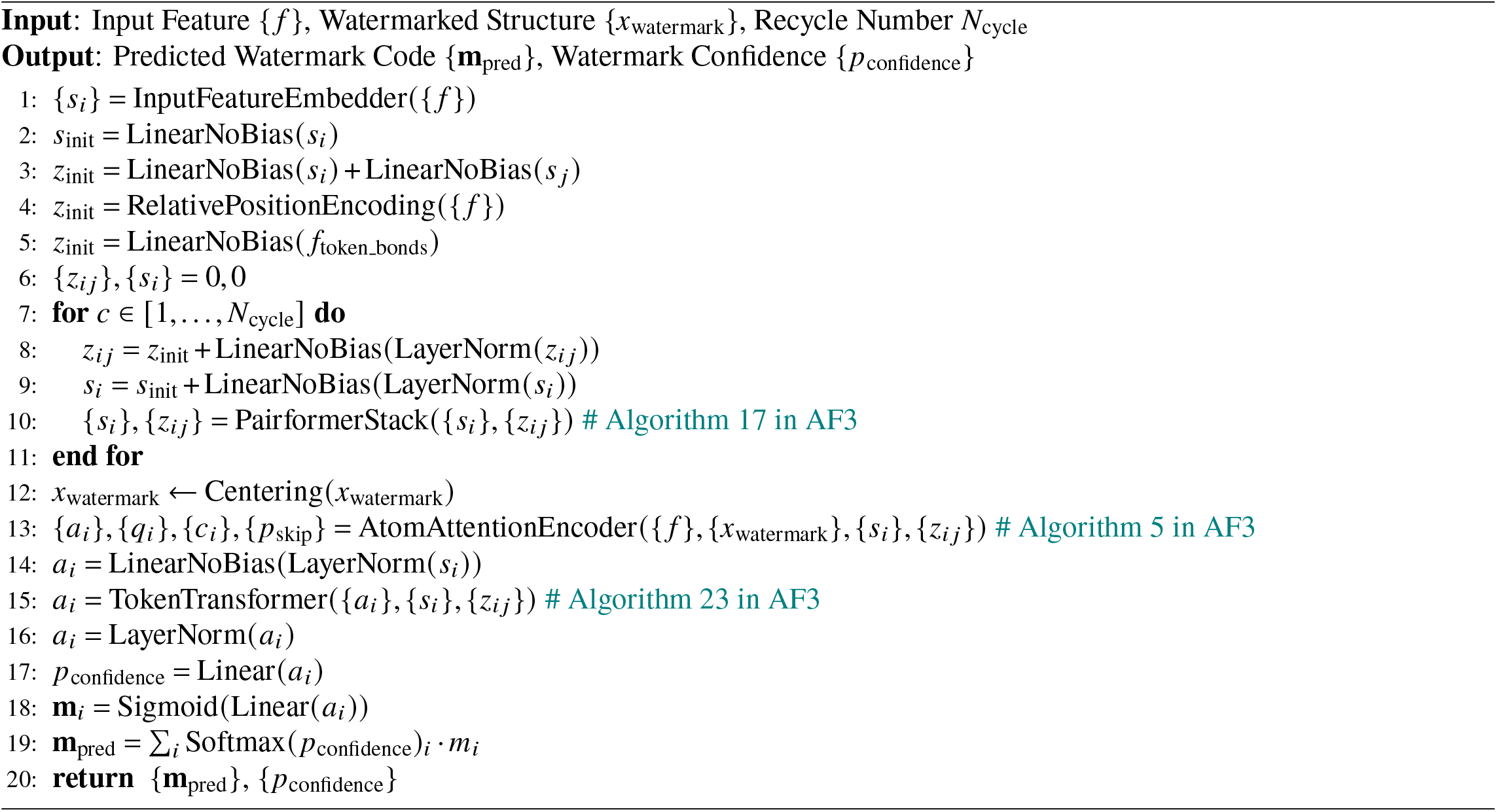

### Introduction and Implementation of Protein Generative Models

#### AlphaFold3

[*3*] (https://github.com/google-deepmind/alphafold3) is the latest non-equivariant diffusion model for biomolecular structures developed by DeepMind, which is capable of predicting the joint structure of complexes including proteins, nucleic acids, small molecules, ions, and modified residues. For MSA, AlphaFold3 uses UniRef90 (v2022 05) [*49*], UniProt (v2021 04) [*80*], Reduced BFD [*81*], MGnify (v2022 05) [*82*] for protein chains and Rfam (v14.9) [*83*], RNACentral (v21.0) [*84*], and Nucleotide collection (2023-02-23) [*85*] for RNA chains. Template structures are retrieved from the corresponding PDB mmCIF file of the searched UniRef90 [*49*] MSA for protein chains. For inference, the number of diffusion steps is 200 and the number of predictions per seed is 5.

#### Protenix

[*54*] (https://github.com/bytedance/Protenix) is an open-sourced version of AlphaFold3, providing model weights, inference code, and trainable code for research use. Generally, Protenix achieves comparable performance with AF3 in predicting structures across different molecular types. For MSA, Protenix searches MSAs using MMseqs2 [*48*] and ColabFold [*67*] MSA pipeline, and use MSAs from the UniRef100 [*49*] database for pairing (species are identified with taxonomy IDs). Protenix does not use MSA for nucleic chains. Protenix also does not use templates. For inference, the number of diffusion steps is 200 and the number of predictions per seed is 5.

#### Boltz-1

[*55*] (https://github.com/jwohlwend/boltz) is an open-source AF3-like model incorporating innovations in model architecture, speed optimization, and data processing, achieving AlphaFold3-level accuracy in predicting the 3D structures of biomolecular complexes. For example, to overcome the potential structure misalignment issue during model inference, Boltz-1 adds a rigid alignment with Kabsch algorithm [*86*] after every step during the inference procedure before 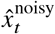 and 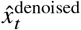 are interpolated. Under the assumption of a Dirac distribution, the interpolated structure is more similar to the denoised sample than the noisy structure. MSAs are constructed for the full PDB data using the colabfold search tool [*67*]. Unlike AF3, Boltz-1 does not include input templates, due to their limited impact on the performance of large models. The training and inference code, model weights, datasets, and benchmarks of Boltz-1 are released under the MIT open license, aiming at democratizing biomolecular interaction modeling.

#### Chai-1

[*56*] (https://github.com/chaidiscovery/chai-lab) is a AF3-like model for molecular structure prediction developed by Chai Discovery. The Chai-1 neural architecture largely follows AF3. One key difference of Chai-1 is the incorporation of protein language model embeddings (ESM-3B) [*45*] as optional input. Per-residue embeddings from ESM are added into single representations of protein chains. Residues that do not belong to a protein chain (e.g., DNA, RNA, or ligands) are assigned a mask token. During inference, Chai-1 can be run with any combination of MSAs, templates, and language model embeddings. MSAs are generated and deposited in OpenProteinSet [*87*] for databases UniRef90 [*49*], UniProt [*80*], MGnify [*82*], and UniClust30+BFD [*81*]. Templates are generated using PDB70 [*50*].

#### ESMFold

[*45*] (https://github.com/facebookresearch/esm) is based on ESM protein language models to generate accurate structure predictions end to end directly from the sequence of a protein. This difference significantly reduces inference time, making ESMFold approximately ten times faster than AlphaFold series. The structure folding head begins with 48 folding blocks. Each folding block alternates between updating a sequence representation and a pairwise representation. The output of these blocks is passed to an equivariant transformer structure module, and 3 steps of recycling are performed before outputting a final atomic-level structure and predicted confidence.

#### RFDiffusion

[*4*] (https://github.com/RosettaCommons/RFdiffusion) is one of the state-of-the-art method for *de novo* protein backbone generation. It combines the RoseTTAFold structure prediction network with the diffusion probabilistic models (DDPMs) framework. When generating ligand-binding protein binders, we use a heuristic attractive-repulsive potential to encourage the formation of pockets with shape complementarity to a target molecule following the suggestions of RFDiffusion [*4*] and RFDiffusionAA [*46*].

#### RFDiffusionAA

[*46*] (https://github.com/baker-laboratory/rf_diffusion_all_atom) is the latest version of RFDiffusion which combines a residue-based representation of amino acids and atomic representations of all other groups to model protein-small molecules/metals/nucleic acids/covalent modification complexes. Starting from random distributions of amino acid residues surrounding target small molecules, RFDiffusionAA can directly generate the small molecule binding protein backbone.

#### FrameDiff

[*51*] (https://github.com/jasonkyuyim/se3_diffusion) is a diffusion model designed for generating protein backbones, leveraging SE(3) symmetry to ensure the generated structures respect the geometric properties of proteins. It operates on frames, which are rigid bodies in 3D space, and employs an SE(3)-invariant diffusion process to create new protein structures, marking a significant advancement in de novo protein design. The model’s architecture includes an SE(3) equivariant score function over multiple frames, developed through a theoretical framework for diffusion on SE(3). This allows FrameDiff to generate designable protein monomers up to 500 amino acids in length without relying on a pre-trained protein structure prediction network.

#### FrameFlow

[*53*] (https://github.com/microsoft/protein-frame-flow) is a method for fast protein backbone generation, utilizing SE(3) flow matching, which adapts the state-of-the-art diffusion model FrameDiff to a flow-matching generative modeling paradigm. This adaptation is significant as it reduces the number of sampling timesteps required. The technical architecture of FrameFlow includes an SE(3)-equivarient neural network, specifically the FramePred architecture with IPA updates. It introduces modifications such as weighting the rotation loss by 0.5, clipping the scaling factor at 1/(1-mint,0.9), and using an exponential scheduler *κ* (*t*) = *e* ^(−*ct*)^ for SO(3) inference, with c=10 found effective.

#### FoldFlow2

[*52*] (https://github.com/DreamFold/FoldFlow) is a state-of-the-art protein structure generative model that integrates both protein sequence and structure information, building on its predecessor, FoldFlow. It employs a multi-modal approach, combining sequence encoding from the ESM-2 protein language model, trained on millions of sequences, with structure encoding from FoldFlow’s structure module. This dual approach addresses the interdependence of sequence and structure, a shortcoming in previous models that tackled them separately. The architecture features a trunk network with multiple attention-based layers for fusing sequence and structure data, ensuring effective integration. The structure decoder mirrors the encoder, providing geometrically informed predictions. FoldFlow2 is trained on a dataset eight times larger than its predecessor FoldFlow. This large-scale training enhances its capability to generate designable, diverse, and novel proteins.

### Supplementary Text

#### More Comparison with Other Watermarking Methods

Traditional watermarking techniques developed for Large Language Models (LLMs) [*33, 88, 89, 90*] and diffusion models [*40, 41, 39, 91*] are not directly transferable to protein structure data due to the distinct and complex characteristics of protein structures. Protein structures exhibit flexibility, sensitivity, and geometric intricacy, requiring specialized methods for embedding and retrieving watermarks without compromising data integrity or model performance.

Previous methods, such as WaDiff [*40*] and AquaLoRA [*41*], embed watermarks into the U-Net backbone of Stable Diffusion models for image generation. While effective in the image domain, these approaches face significant challenges in protein generative models. The use of convolutional neural networks (CNNs) as encoder-decoder components in WaDiff and AquaLoRA limits their performance in the protein domain, as CNNs are not inherently designed to handle the spatial and rotational properties of protein structures. FoldMark overcomes these limitations by leveraging state-of-the-art SE(3)-equivariant or non-equivariant graph transformers for both the Watermark Encoder and Decoder, ensuring geometric consistency and superior performance. Additionally, FoldMark introduces customized data augmentation strategies, such as structure cropping, rotation, and noising, to enhance the robustness of training. These strategies are tailored to protein structures and are not directly transferable from image-based methods. To address the challenges of protein flexibility and sensitivity, FoldMark adaptively embed watermark information across all residues guided by evolutionary signals (e.g., genetic search, protein language model embeddings, and structure templates). Therefore, FoldMarks learsn to minimize noise in conserved regions while exploiting flexible regions for enhanced watermark capacity.

FoldMark employs a gating vector derived from the watermark code for watermark-conditioned Low-Rank Adaptation (LoRA), ensuring independence from rank selection. This design avoids the parameter overhead associated with AquaLoRA’s reliance on diagonal matrix modifications, which require large ranks (e.g., 320) to embed watermark information effectively.

Finally, FoldMark incorporates an additional, time-conditioned fine-tuning step for the watermark decoder, allowing it to adapt to the intricate protein structures. The message retrieval loss assigns dynamic weights to different time steps (e.g., larger weights as *t* →0). This approach balances the preservation of generation quality with explicit improvements in watermark retrieval success rates.

In summary, FoldMark addresses the unique challenges of protein generative models by combining advanced architectural design, robust watermark strategies, and innovative fine-tuning approaches, achieving superior performance in protecting protein generative models compared to existing methods.

#### Influence of FoldMark on Inverse Folding

**Figure S1.**
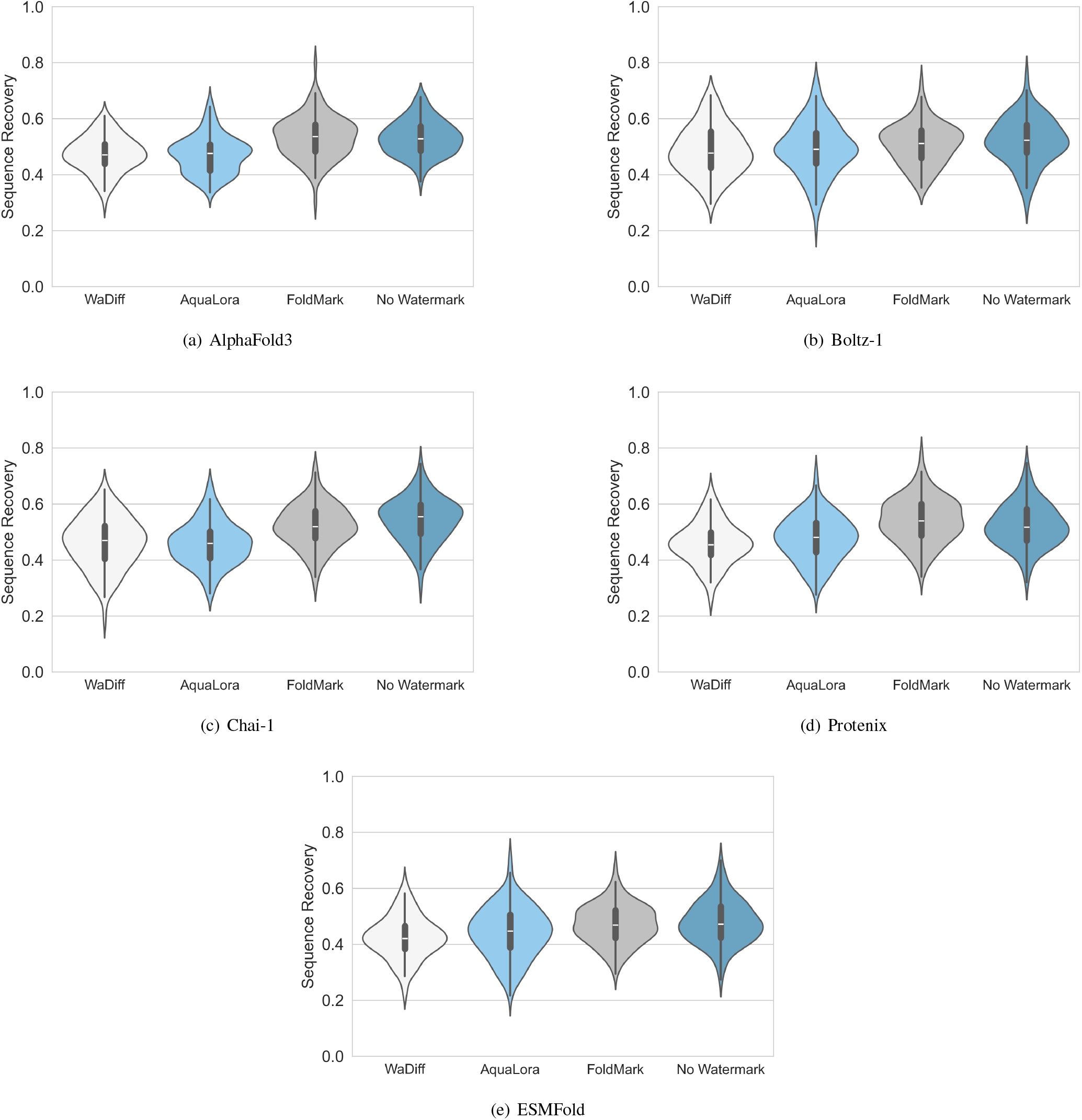
Sequence Recovery Performance of Predicted Structures Generated by WaDiff, AquaLora, FoldMark, and Baseline Vanilla Models. Experiments were conducted on filtered single-chain protein datasets with AlphaFold3, Boltz-1, Chai-1, Protenix, and ESMFold models. Protein sequences were inferred using ProteinMPNN from the predicted structures, and sequence recovery rates were calculated as the percentage of correctly predicted amino acids relative to the native sequences, highlighting the models’ ability to recapitulate biologically relevant protein sequences. In the violin plots, the embedded box plot includes a thick vertical bar indicates the interquartile range (IQR, 25th to 75th percentiles) and a horizontal line marking the median. Thin whiskers extend to the minimum and maximum values within 1.5 times the IQR.

#### Ablation Studies of FoldMark

**Figure S2.**
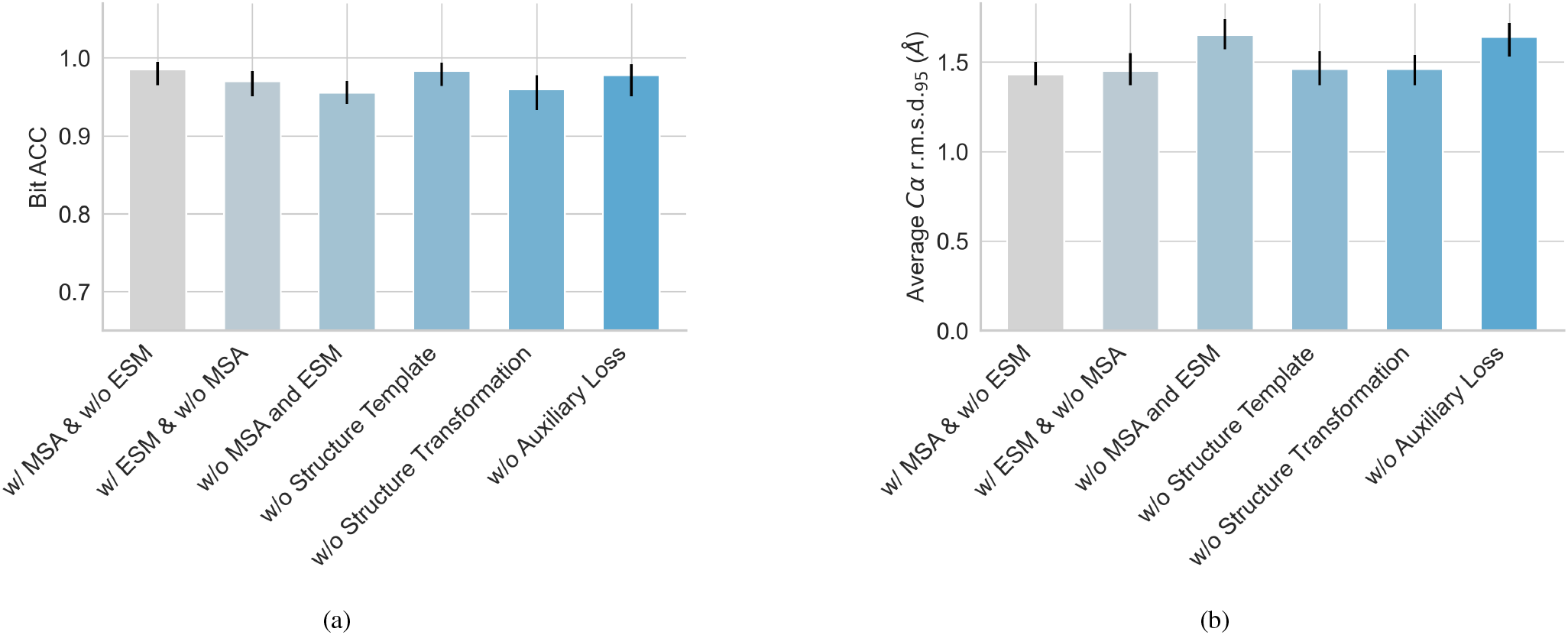
Ablation studies of FoldMark with AF3. The MSA, ESM, and Template modules in FoldMark Encoder, the structure transformations in FoldMark Encoder/Decoder pertaining, and the auxiliary loss in FoldMark Finetuning are removed respectively. Note that the FoldMark Encoder can utilize either the MSA module, the ESM module, or operate without any MSA/ESM module. The shadows indicate the 95% confidence intervals obtained by 10,000 bootstrap resampling.

#### Hyperparameter Analysis of FoldMark

**Figure S3.**
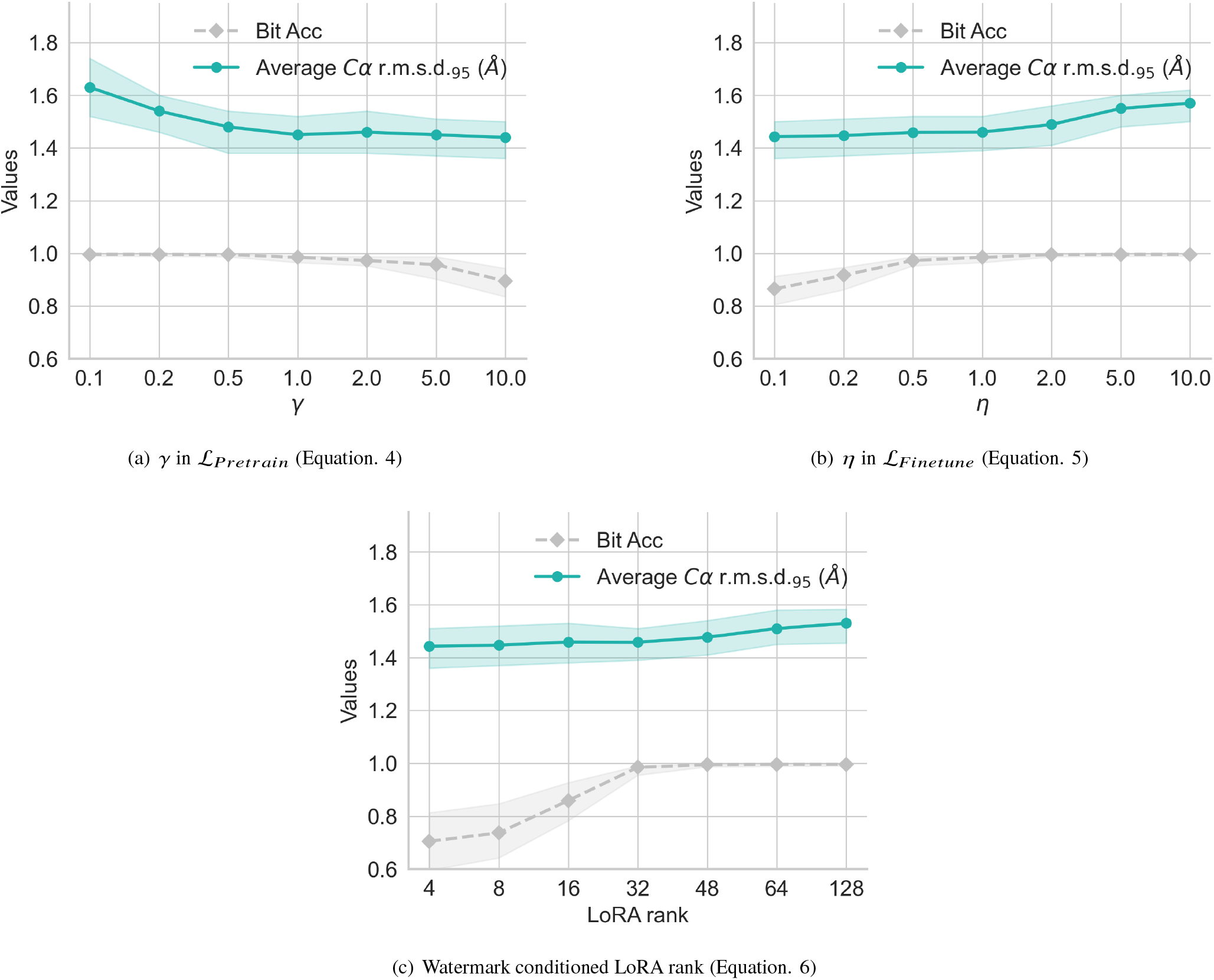
Hyperparameter Analysis of FoldMark. The shadows indicate the 95% confidence intervals obtained by 10,000 bootstrap resampling.

#### More Results on Detecting Unauthorized Use of Watermarked Data

**Figure S4.**
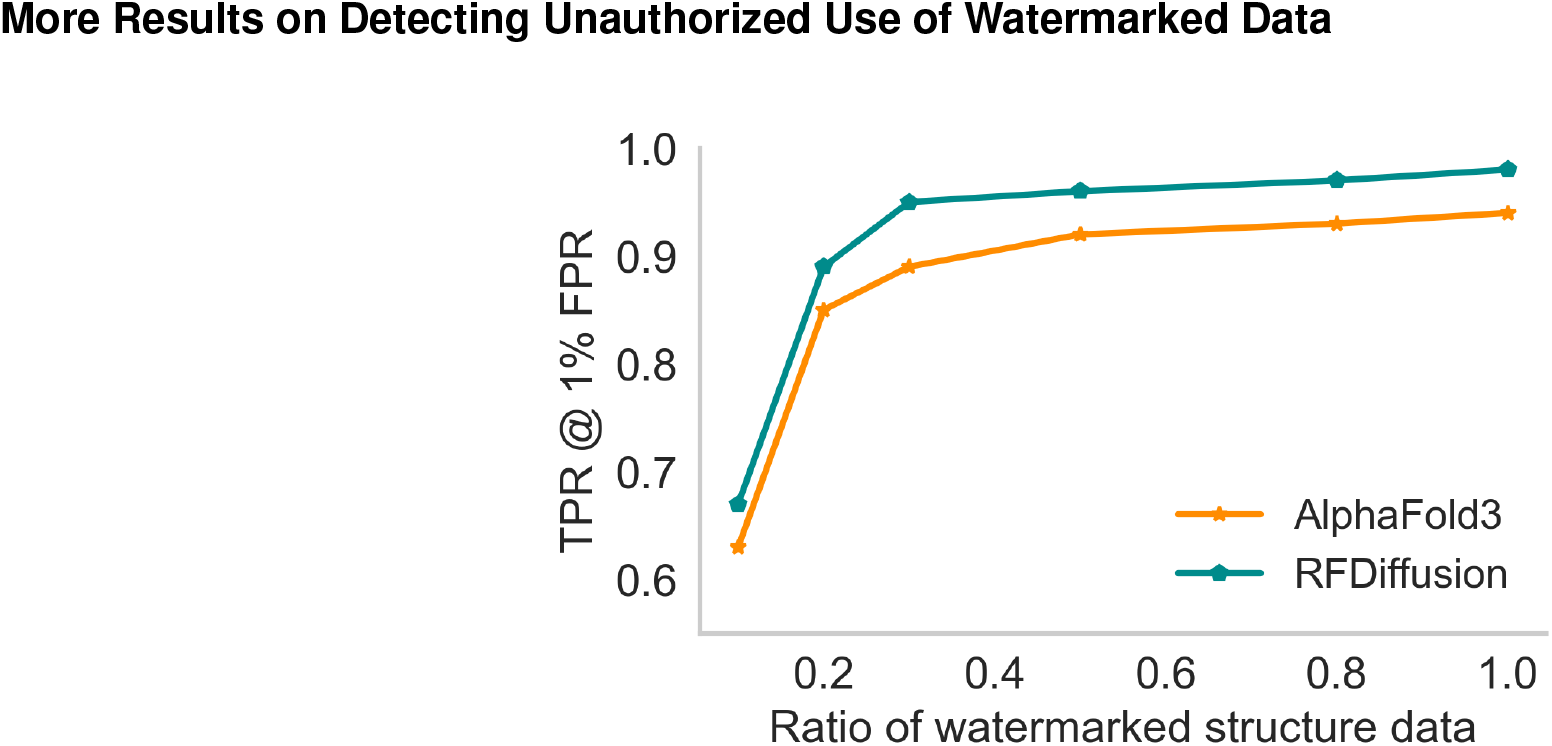
Influence of the ratio of watermarked structures on detection true positive rates at 1% false positive rates.

#### Performance of FoldMark for User Tracing under Different Post-processing

**Figure S5.**
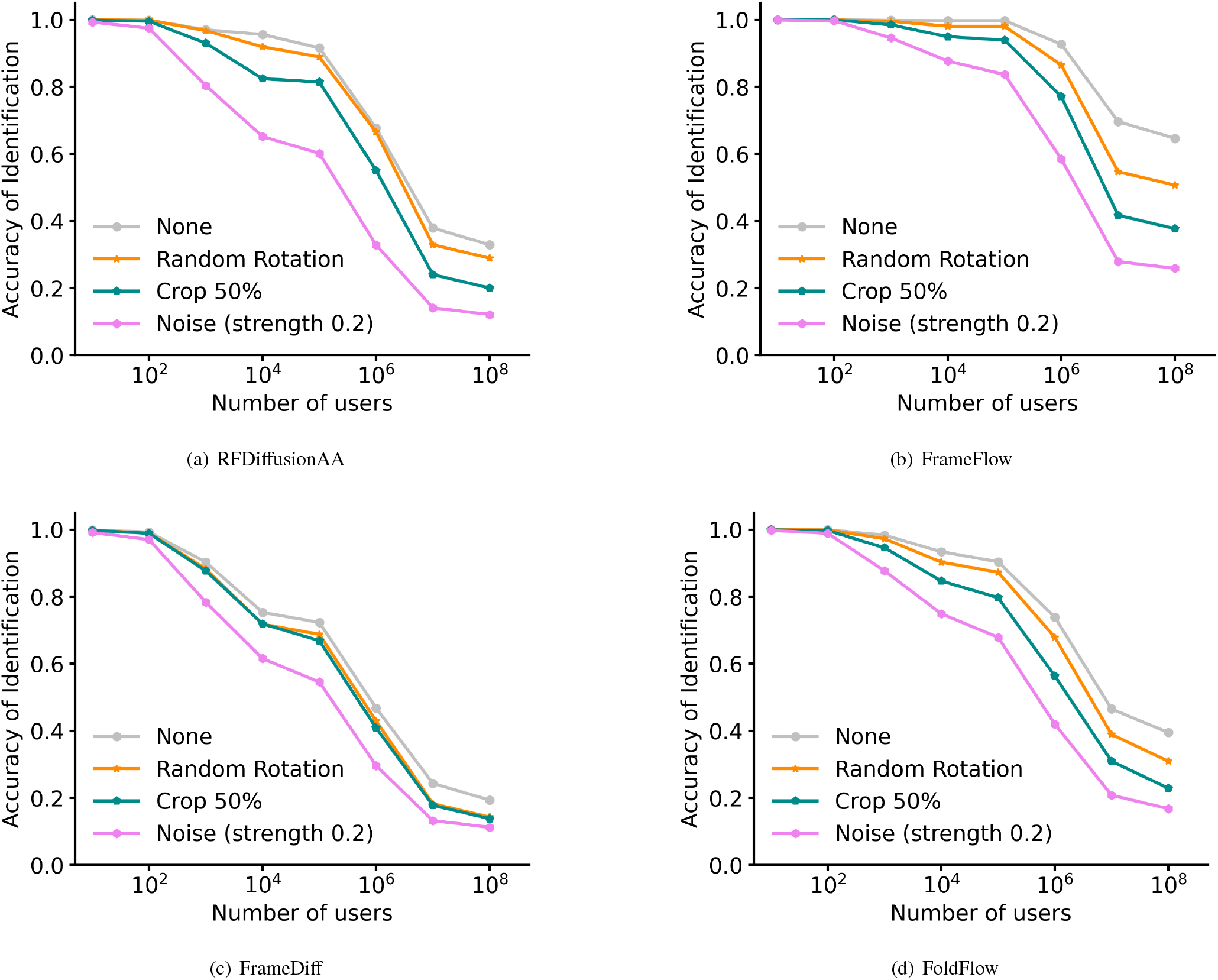
Performance of FoldMark for user tracing under different post-processing with different numbers of users (FPR=10^−3^). (a) RFDiffusionAA; (b) FrameFlow; (c) FrameDiff; (d) FoldFlow are compared here. Protein post-processing methods include randomly rotating the watermarked strictures (Random Rotation), cropping the structure to keep 50% of the whole sequence (Crop 50%), and adding Gaussian noise with a strength of 0.2 to the coordinates (Noise, strength 0.2).

#### Empirical calibration of the null-bit distribution

**Figure S6.**
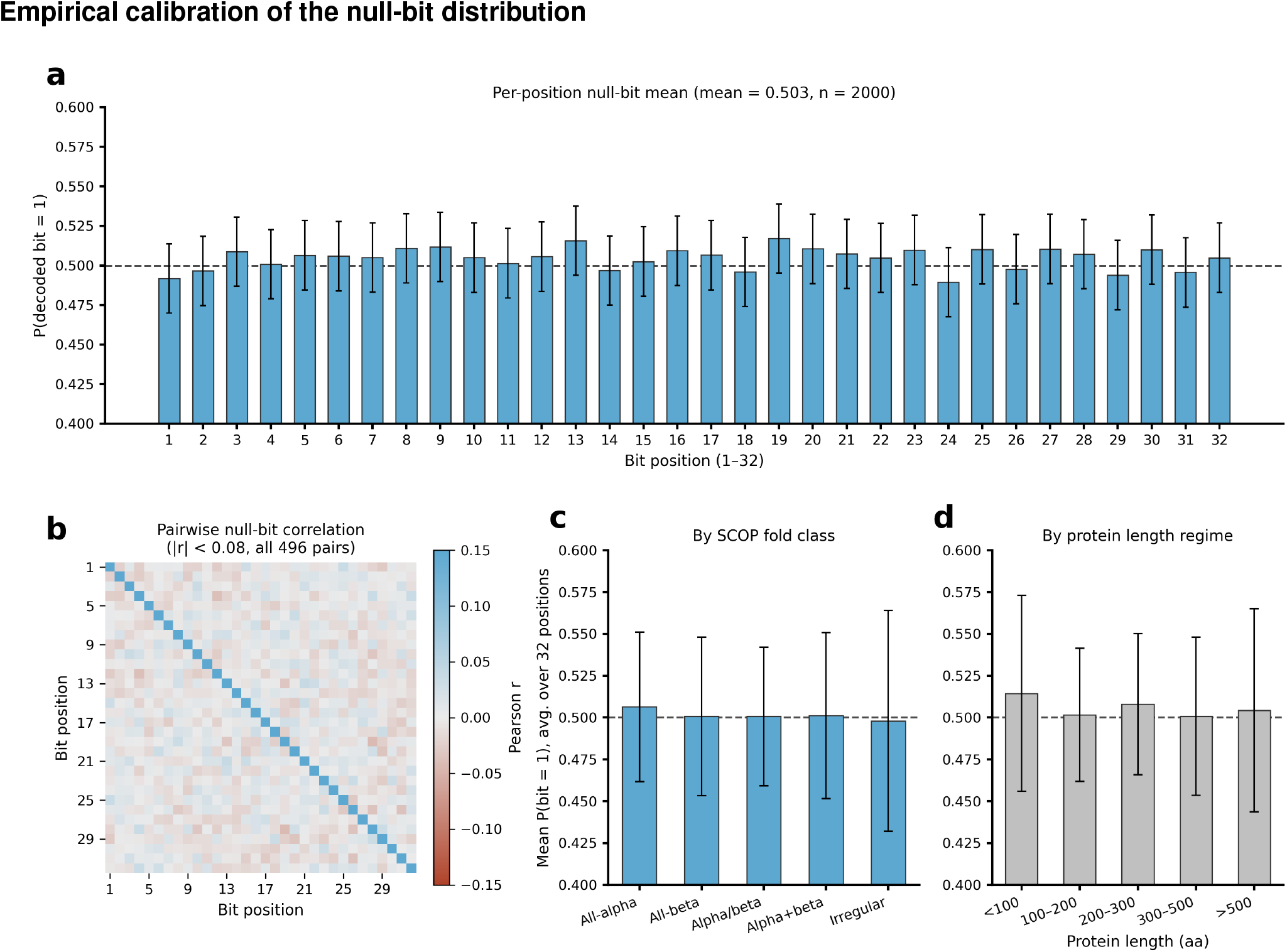
Calibration of decoded bits on unwatermarked structures under the null (no watermark embedded) condition. **(a)** Per-position mean probability of decoding a 1, 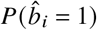, across all 32 code positions (*n* = 2,000 unwatermarked structures; error bars are 95% binomial confidence intervals; dashed line is the ideal value of 0.5). **(b)** Pairwise Pearson correlation matrix between decoded bit positions; all 496 off-diagonal pairs satisfy |*r*| < 0.08, indicating negligible inter-bit correlation. **(c)** Mean 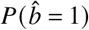, averaged over the 32 positions, stratified by SCOP fold class (all-alpha, all-beta, alpha/beta, alpha+beta, irregular). **(d)** Mean 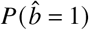 stratified by protein length regime (<100 to >500 residues). Across all strata, the decoded null-bit probability remains close to the ideal value of 0.5, supporting the independent Bernoulli(0.5) null assumption underlying our false-positive calibration and tracing formula.

#### Gating Strategy of Flow Cytometry

**Figure S7.**
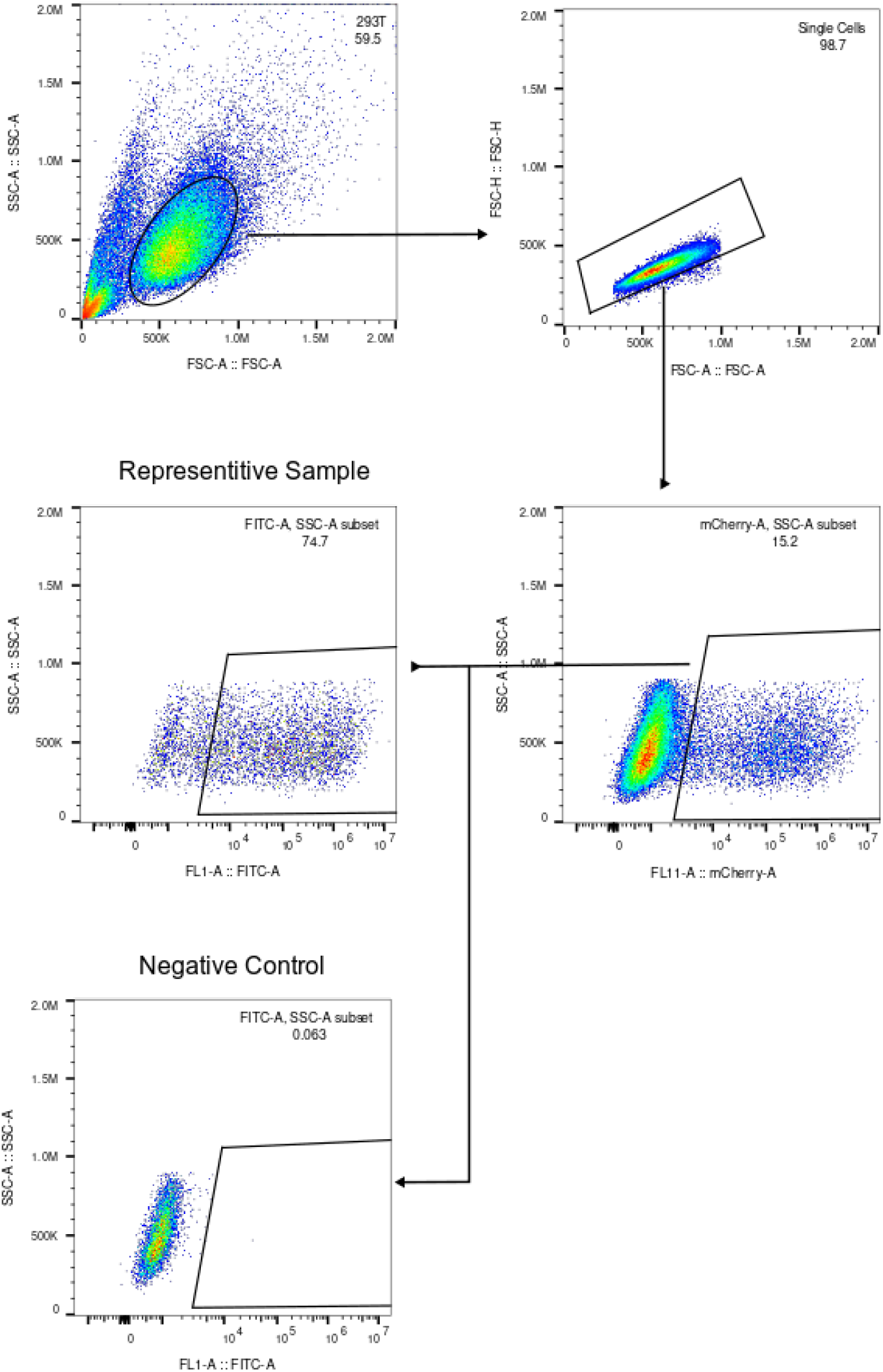
Gating strategy for Cas13bt3 reporter assay.

**Figure S8.**
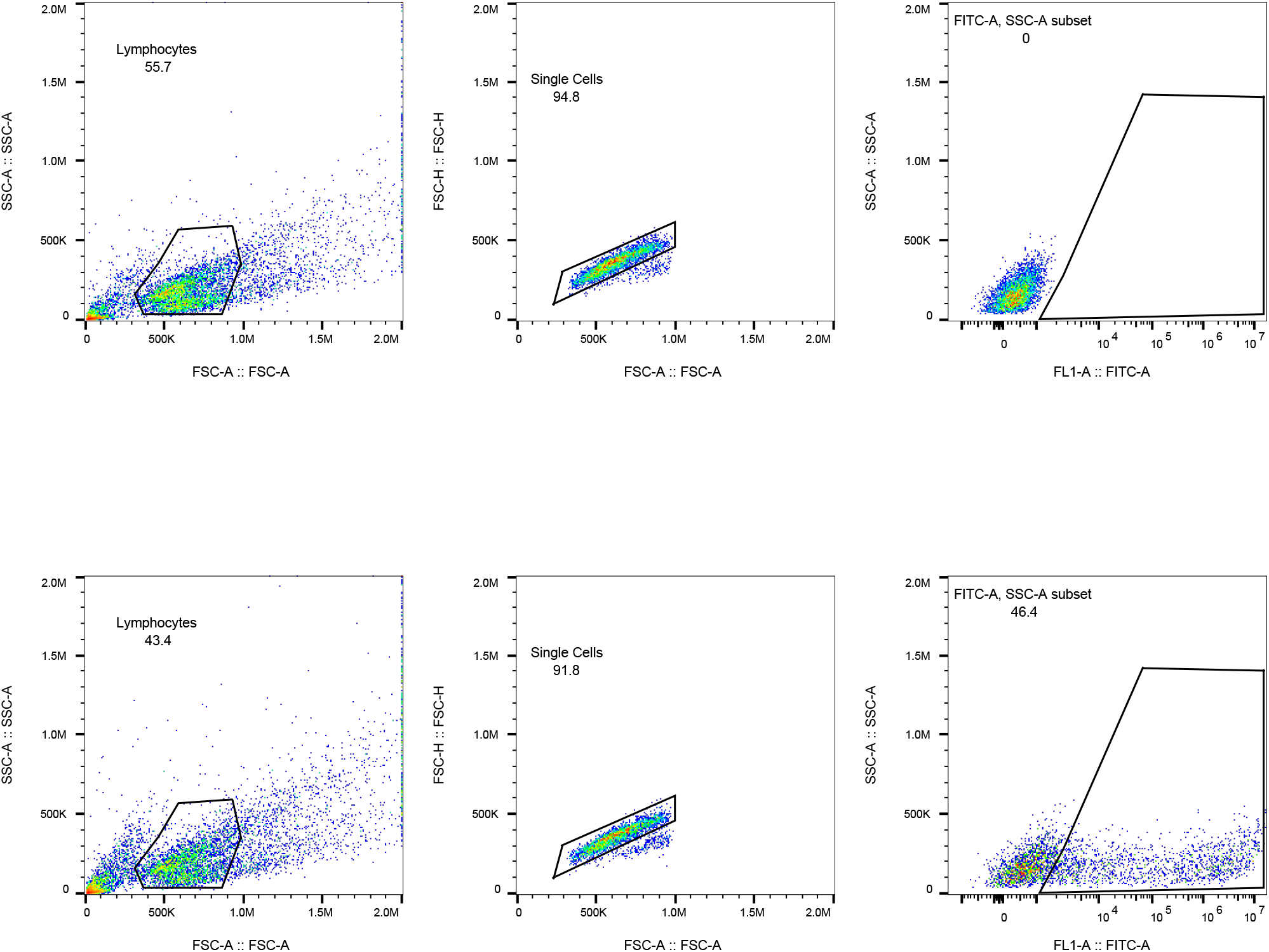
Gating strategy for EGFP flow cytometry analysis. Rows represent the negative control (top) and the EGFP-transfected sample (bottom).

**Figure S9.**
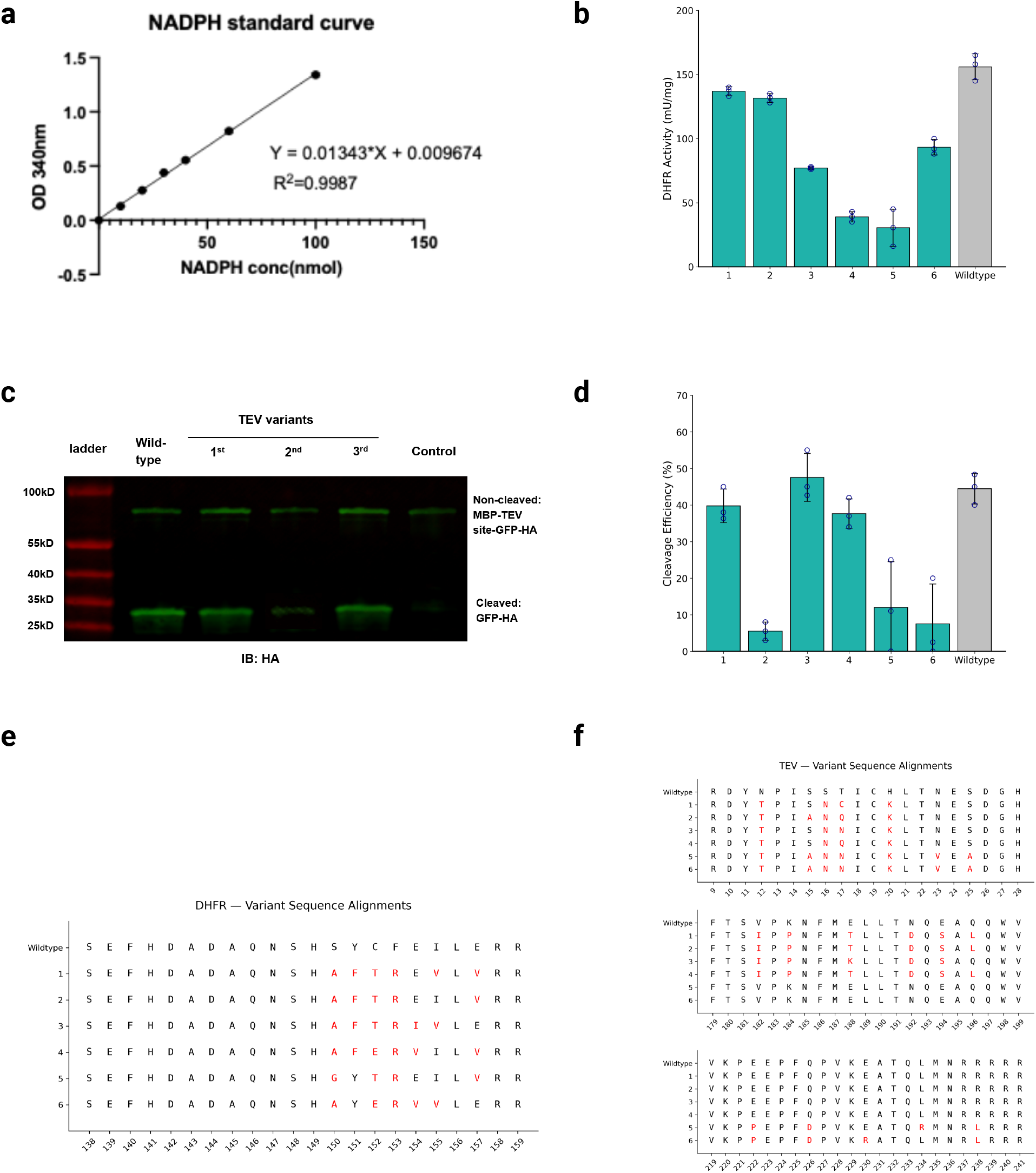
Wet-lab validation of FoldMark-redesigned DHFR and TEV variants. For each enzyme, we generated six FoldMark-watermarked variants and ranked them using ESM-based sequence scoring. The top three variants are shown in the main text, whereas all six tested variants are shown here. **a**, NADPH standard curve used for quantifying DHFR enzymatic activity in the colorimetric assay. The linear regression equation and coefficient of determination are shown. **b**, DHFR activity of all six FoldMark-redesigned DHFR variants compared with wild-type DHFR. Activity was quantified using the NADPH-based colorimetric DHFR assay. Variants are ordered according to their ESM-based ranking. **c**, Representative western blot validating TEV-mediated substrate cleavage. TEV variants were incubated with the MBP–TEV site–GFP–HA substrate, and the non-cleaved substrate and cleaved GFP–HA product were detected by immunoblotting with an anti-HA antibody (IB: HA). **d**, Quantification of TEV cleavage efficiency for all six FoldMark-redesigned TEV variants compared with wild-type TEV. Cleavage efficiency was calculated from the intensities of the cleaved GFP–HA product and non-cleaved MBP–TEV site–GFP–HA substrate. Variants are ordered according to their ESM-based ranking. **e**, Sequence alignment of wild-type DHFR and all six FoldMark-redesigned DHFR variants. Watermark-associated mutation sites are highlighted in red. **f**, Sequence alignment of wild-type TEV and all six FoldMark-redesigned TEV variants. Watermark-associated mutation sites are highlighted in red. **47/50**

#### More Results of FoldMark for Different Models

**Table S1.** Watermarking different protein structure prediction models by FoldMark with different watermark bits. The average watermark bit accuracy (BitAcc) and the Cα r.m.s.d. at 95% coverage (RMSD) are reported.

| Watermark Method |  | 4bit |  | 8bit |  | 16bit |  | 32bit |  |
| --- | --- | --- | --- | --- | --- | --- | --- | --- | --- |
| | | BitAcc ↑ | RMSD ↓ | BitAcc ↑ | RMSD ↓ | BitAcc ↑ | RMSD ↓ | BitAcc ↑ | RMSD ↓ |
| Protenix | No watermark | - | 1.464 | - | 1.464 | - | 1.464 | - | 1.464 |
|  | WaDiff | 66.3% | 1.542 | 64.0% | 1.583 | 60.4% | 1.606 | 52.1% | 1.723 |
|  | AquaLoRA | 64.0% | 1.526 | 62.9% | 1.530 | 61.2% | 1.648 | 50.4% | 1.691 |
|  | FoldMark(Ours) | **99.5%** | **1.465** | **98.5%** | **1.468** | **97.3%** | **1.470** | **95.4%** | **1.568** |
| ESMFold | No watermark | - | 1.807 | - | 1.807 | - | 1.807 | - | 1.807 |
|  | WaDiff | 78.5% | 1.920 | 74.1% | 1.943 | 71.6% | 2.069 | 54.0% | 2.200 |
|  | AquaLoRA | 79.4% | 1.895 | 76.8% | 1.957 | 75.3% | 2.055 | 59.4% | 2.120 |
|  | FoldMark(Ours) | **99.6%** | **1.815** | **97.2%** | **1.819** | **96.2%** | **1.821** | **90.1%** | **1.845** |

**Table S2.** Watermarking *de novo* protein structure generative models by FoldMark with different watermark bits. The average watermark bit accuracy (BitAcc) and the self-consistency TM scores (scTM) are reported.

| Watermark Method |  | 4bit |  | 8bit |  | 16bit |  | 32bit |  |
| --- | --- | --- | --- | --- | --- | --- | --- | --- | --- |
| | | BitAcc ↑ | scTM ↑ | BitAcc ↑ | scTM ↑ | BitAcc ↑ | scTM ↑ | BitAcc ↑ | scTM ↑ |
| FrameFlow | No watermark | - | 0.921 | - | 0.921 | - | 0.921 | - | 0.921 |
|  | WaDiff | 77.1% | 0.875 | 76.4% | 0.836 | 70.5% | 0.826 | 54.6% | 0.811 |
|  | AquaLoRA | 79.6% | 0.860 | 75.4% | 0.829 | 74.5% | 0.810 | 55.1% | 0.805 |
|  | FoldMark(Ours) | **99.6%** | **0.920** | **99.5%** | **0.919** | **96.7%** | **0.916** | **95.4%** | **0.904** |
| FrameDiff | No watermark | - | 0.890 | - | 0.890 | - | 0.890 | - | 0.890 |
|  | WaDiff | 76.8% | 0.863 | 73.3% | 0.848 | 62.2% | 0.810 | 50.4% | 0.795 |
|  | AquaLoRA | 64.3% | 0.852 | 59.1% | 0.839 | 56.2% | 0.794 | 51.6% | 0.778 |
|  | FoldMark(Ours) | **98.7%** | **0.890** | **98.3%** | **0.875** | **88.4%** | **0.873** | **82.0%** | **0.850** |
| FoldFlow | No watermark | - | 0.945 | - | 0.945 | - | 0.945 | - | 0.945 |
|  | WaDiff | 88.2% | 0.869 | 85.0% | 0.828 | 80.1% | 0.777 | 64.3% | 0.773 |
|  | AquaLoRA | 73.5% | 0.854 | 72.6% | 0.830 | 71.7% | 0.786 | 62.8% | 0.778 |
|  | FoldMark(Ours) | **99.9%** | **0.937** | **99.7%** | **0.936** | **98.9%** | **0.931** | **94.5%** | **0.924** |

**Table S3.** Watermark detectability after independent clean forward structure prediction. For each protein, the positive set consisted of watermarked redesigned sequences generated from the FoldMark-watermarked backbone. The negative set consisted of *matched non-watermarked pipeline controls*, rather than unmodified wildtype sequences. Specifically, the negative controls were generated from the corresponding non-watermarked backbone using the same redesigned regions, inverse-folding procedure, candidate-pool size, mutation-count constraints, mutation caps, and ESM-based candidate-ranking procedure as the positive set. Both sets were subsequently folded using independent clean structure predictors that were not involved in the FoldMark inverse-folding or refolding pipeline. Watermark classification was evaluated in a balanced binary setting with equal numbers of watermarked sequences and matched non-watermarked pipeline controls. TPR denotes the fraction of watermarked sequences correctly detected, whereas FPR denotes the fraction of matched non-watermarked controls incorrectly classified as watermarked.

| Protein | Clean forward predictor | Accuracy (%) | TPR (%) | FPR (%) | F1 score (%) |
| --- | --- | --- | --- | --- | --- |
| EGFP | AlphaFold3 | 99.0 | 100.0 | 2.0 | 99.0 |
| EGFP | ESMFold | 96.0 | 96.0 | 4.0 | 96.0 |
| EGFP | Protenix | 98.0 | 98.0 | 2.0 | 98.0 |
| EGFP | Boltz-1 | 97.0 | 98.0 | 4.0 | 97.0 |
| EGFP | Chai-1 | 100.0 | 100.0 | 0.0 | 100.0 |
| Cas13 | AlphaFold3 | 97.0 | 96.0 | 2.0 | 97.0 |
| Cas13 | ESMFold | 95.0 | 94.0 | 4.0 | 94.9 |
| Cas13 | Protenix | 99.0 | 100.0 | 2.0 | 99.0 |
| Cas13 | Boltz-1 | 98.0 | 98.0 | 2.0 | 98.0 |
| Cas13 | Chai-1 | 98.0 | 98.0 | 2.0 | 98.0 |
| **Average** |  | **97.7** | **97.8** | **2.4** | **97.7** |

**Table S4.**
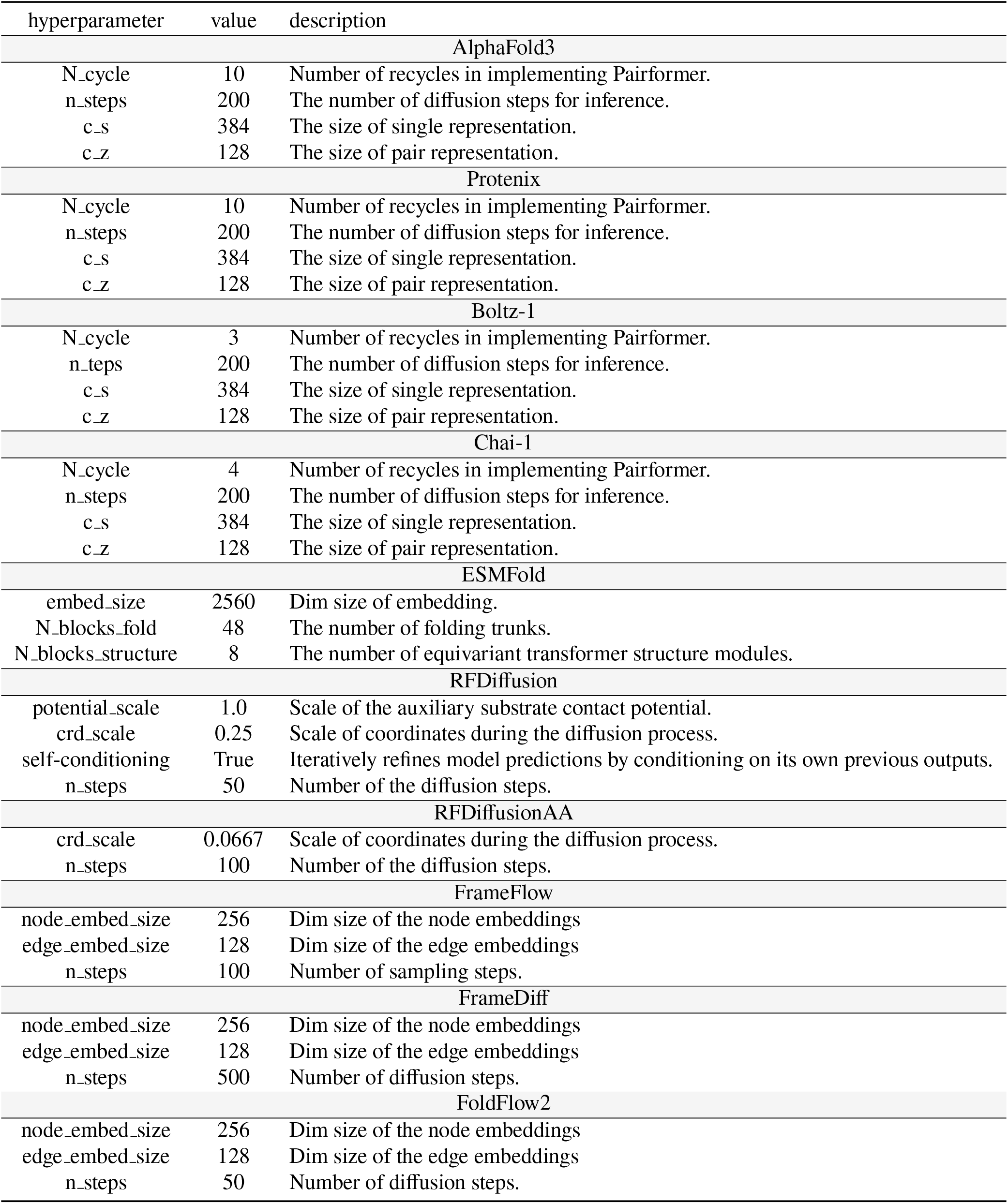
Hyperparameters for protein generative models.

In Table. S1 and S2, we show more detailed results of applying FoldMark to different protein structure prediction models and *de novo* protein structure generative models.

## Footnotes

1 https://github.com/rmin2000/WaDiff

2 https://github.com/Georgefwt/AquaLoRA

